# MYXOBACTERIA FROM VIETNAM: ISOLATION, PHYLOGENETIC ANALYSIS AND BIOPROSPECTION

**DOI:** 10.1101/2022.04.27.489816

**Authors:** Yen Thi Ngoc Nguyen, Chung Dinh Duong, Hong Thi Kim Nguyen, Cham Phuong Nguyen, Nhan Thi Vo, Linh Thi Lan Dinh, Ngoc Le Bao Nguyen, Thai Minh Nguyen, Nga Dinh Nguyen, Anh Tu Nguyen

## Abstract

Forty-three myxobacterial strains were isolated and identified from 20 soil samples collected in Vietnam. The information on morphological characteristics and 16S ribosomal gene sequencing showed that these strains were designated to seven genera belonging to Angiococcus, Archangium, Chondromyces, Corallococcus, Cystobacter, Melittangium, and Myxococcus, in suborder Cystobacterineae and Sorangiineae. The phylogenetic tree was constructed to clarify the genetic relationship between myxobacterial isolates. Myxobacteria were cultured, and crude extracts were obtained after 10-day fermentation in P-medium in the presence of the Amberlite XAD 16N adsorbent resin. Elution was carried out with acetone and methanol to obtain the crude extracts. Evaluation of antioxidant activity used the DPPH and ABTS assay, the minimum inhibitory concentration values were determined by the microdilution method. The total extract from CT21 had the highest total antioxidant activity (IC_50_ = 52.34 ± 1.47 µg/mL, 30.28 ± 0.74 µg/mL for DPPH and ABTS assays, respectively). The other potential strain was TG131 and GL41 that IC_50_ values were 40.28 ± 1.13 and 57.24 ± 1.52 µg/mL, respectively (by the DPPH method), and 48.35 ± 0.58 and 42.76 ± 0.50 µg/mL, respectively (by the ABTS method). Interestingly, 100% isolated myxobacterial strains show inhibitory activity against at least one of the tested microorganisms. The potential antimicrobial strain was GL41, which inhibited all tested microorganisms, and the MIC values were 1 µg/mL against MRSA, MSSA, *S. faecalis, C. albicans*, and *A. niger*. The highest active strains were members of Myxococcus sp. genus.

## 1. Introduction

Myxobacteria is a group of Gram-negative bacteria, long rods, and widespread in natural habitats such as soil, water, bark, herbivore dung, desert, even marine areas, saline-alkaline soil [1]. The adverse conditions facilitate exceptional ability to produce diverse secondary metabolites [2]. Myxobacteria have been documented as a promising microorganism with broad-ranging bioactivities including antioxidant, antivirus, antimicrobial, antimalarial, and cytotoxic activities [3]. More than 100 metabolites and 600 derivatives have been found [4]. So most compounds were novel, not found in other microbes, and exhibited many remarkable modes of action that myxobacteria are regarded as outstanding secondary metabolite producers for drug development [5].

In recent years, many notable compounds continue to be discovered, such as archazolids (*Archangium gephyra*, 2003) [6], chondrochloren (*Chondromyces crocatus*, 2003) [7], leupyrrins (Sorangium cellulosum, 2003) [8], ajudazols (*Chondromyces crocatus*, 2004) [9], aurafuron (*Stigmatella aurantiaca* and *Archangium gephyra*, 2005) [10], cruentaren (*Byssovorax cruenta*, 2006) [11], bithiazole (Myxococcus fulvus, 2007) [12], spirodienal (*Sorangium cellulosum*, 2009) [13], myxoprincomide (*Myxococcus xanthus*, 2011) [14], indiacens A and B (*Sandaracinus amylolyticus*, 2012) [15], salimyxin B and enhygrolide A (*Enhygromyxa salina*, 2013) [16], cystodienoic acid (Cystobacter sp., 2015) [17], pyxipyrrolones (Pyxidicoccus, 2017) [18].

Nevertheless, scientists have still believed that the potential bioactive molecules from myxobacteria have not been fully discovered since the biodiversity in myxobacteria and their products may depend on their different habitats [19].

In this study, we isolated and identified myxobacteria from the soil of some provinces in Vietnam and screened antioxidant and antimicrobial activity from their raw extracts. This is the first study on the isolation, determination of phylogenetic relationships and screening bioactivities of myxobacterial strains conducted in Vietnam.

## 2. Materials and methods

### 2.1. Soil

Soil was collected from 20 provinces in Vietnam at a depth of 10-cm into sterile falcons or bags (Table 1). Then the samples were air-dried as soon as possible to reduce the natural moisture and inhibit the development of fungi and worms [20].

**Table 1.**
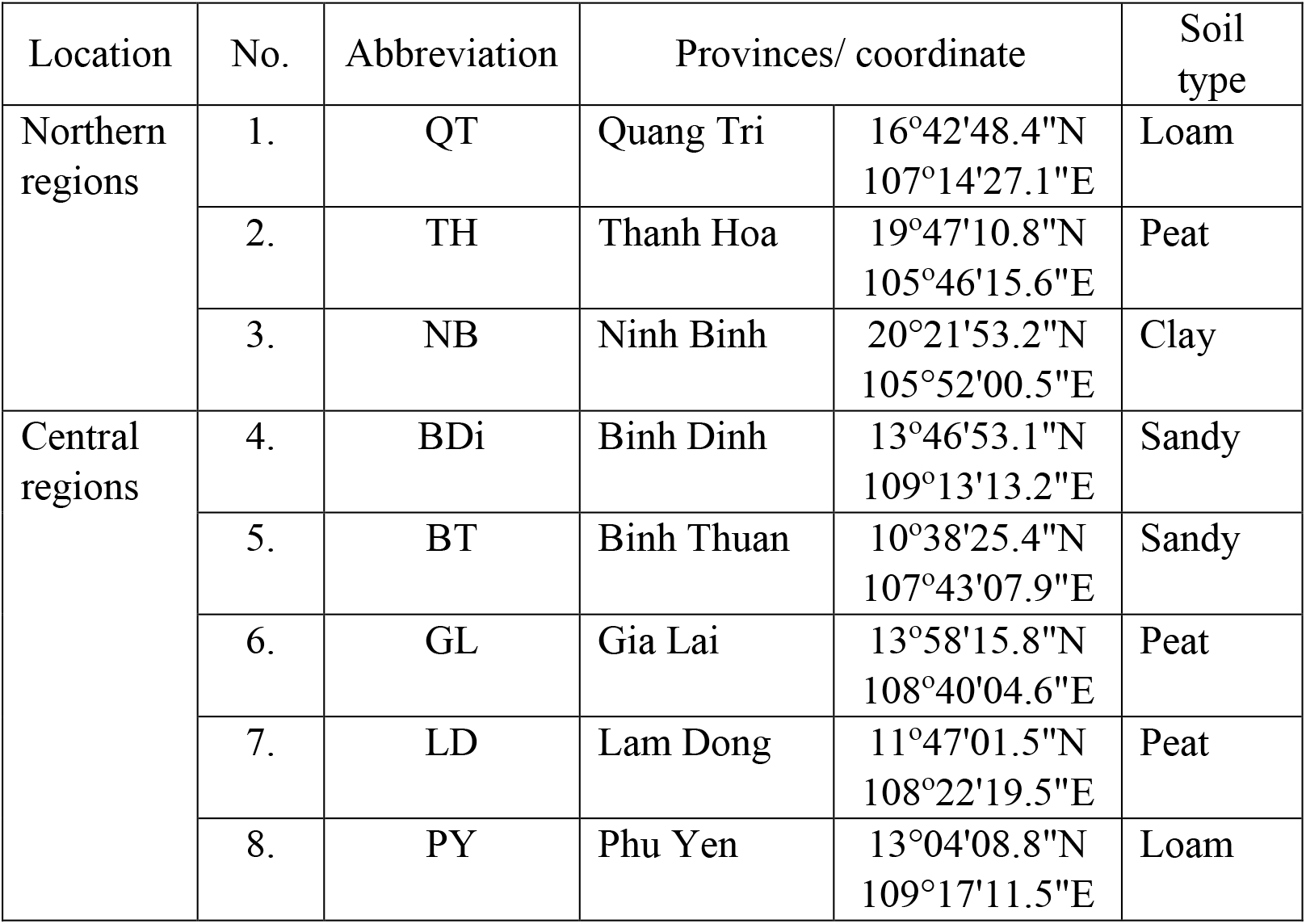
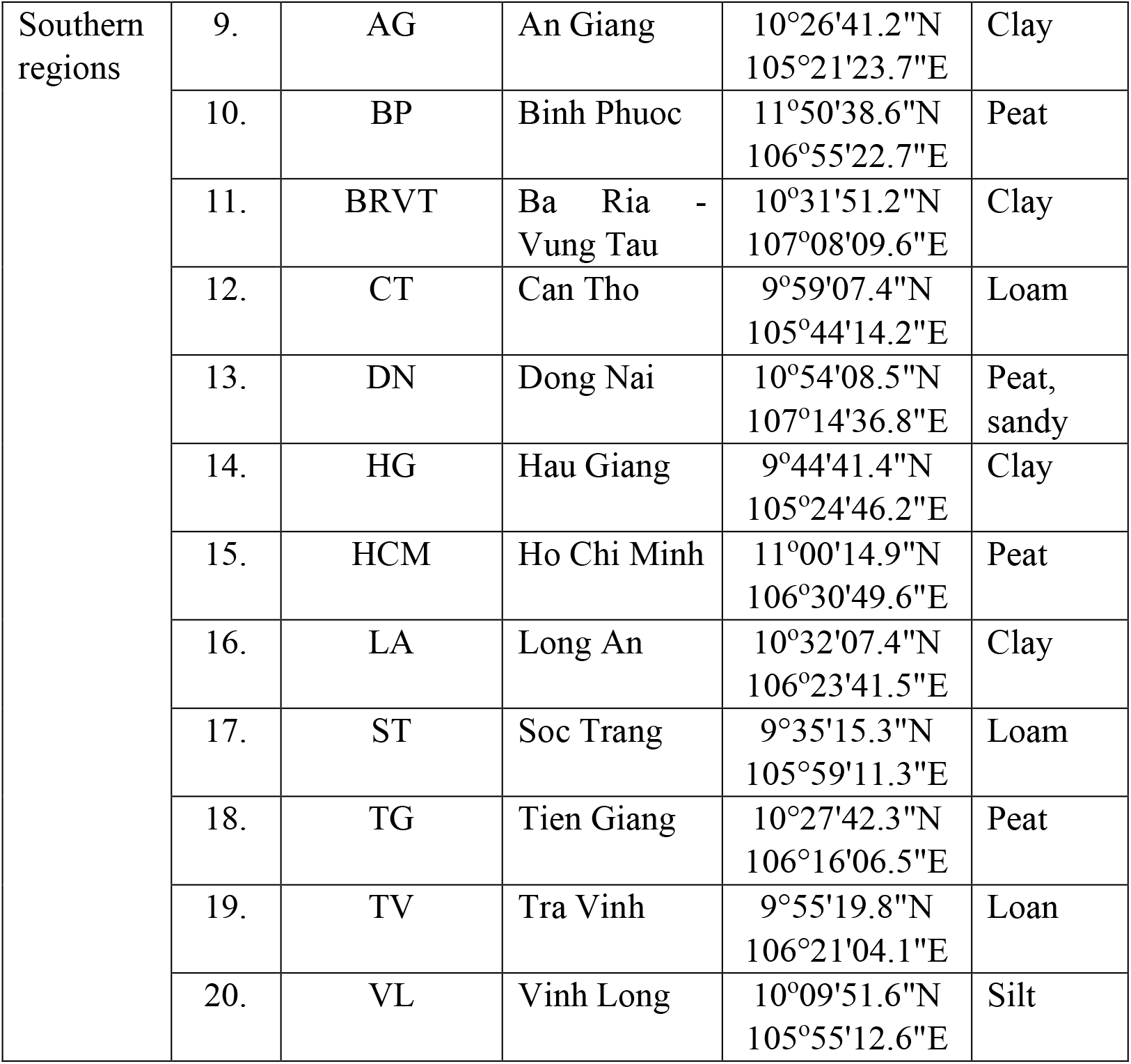
Sampling location in Vietnam

**Table 2.** Morphological characteristics of 5 genera

|  | *Archangium* | *Corallococcus* | *Chondromyces* | *Cystobacter* | *Myxococcus* |
| --- | --- | --- | --- | --- | --- |
| Fruiting body | The large and stretched cluster of sporangioles | Relatively small, various shapes, sporadic scattered on the agar surface | Long, unbranched, or branched stalks; clusters of myxospores attached to the top of the stalk | Cluster or chained, oval, brown or dark pink | Globular or oval, solitary or masses, yellow, pink, red or brown |
| Colony | Covered by slime sheet with radial lines | Large swarm, easy to peel off from the surface | Broad, shallow on the agar surface, with a band of orange cells at the edge | Large colony, covered by slime layer with radial lines | Large, transparent colony and lobes edge |

### 2.2. Isolation and purification of the myxobacteria

Myxobacteria were isolated from ground soil based on three typical methods including (i) WCX medium (w/v, agar 1.5%, CaCl_2_ 0.07%, cycloheximide 0.01%) with *E. coli* streaks (*E. coli* was cultivated in Luria-Bertani broth) for bacteriolytic strains (W), (ii) ST21CX medium (w/v, agar 1%, cycloheximide 0.01%, CaCl_2_.2H_2_O 0.1%, FeCl_3_ 0.02%, K_2_HPO_4_ 0.1%, KNO_3_0.1%, MgSO_4_.7H_2_O 0.1%, MnSO_4_.7H_2_O 0.01%, yeast extract 0.002%) with filter paper for cellulolytic decomposers (F), and (iii) wild rabbit dung method that contain natural substrate suitable for Myxococcus and Corallococcus isolation (R) [20].

Dried soil was heated in a water bath at 80°C for 30 minutes before filling half the Petri dish. Cycloheximide solution 100 µg/mL and amphotericin B 10 µg/mL were added to reduce the overwhelming of fungi. The Petri dishes were wrapped with paraffin, incubated at 30°C for 3-30 days, and checked by stereoscopy daily to recognize the fruiting bodies’ appearances. Myxobacterial isolates were purified by transferring the swarm edge and fruiting bodies many times on VY2 dishes. The purity of isolates was examed by the CEH medium for 24 h [20].

### 2.3. Bacterial identification

#### 2.3.1. Morphology

Isolated bacteria were observed in their colonies, size and shape after Gram staining, type of fruiting bodies and compared with The Prokaryote: Deltaproteobacteria and Epsilonproteobacteria [21] and Order VIII. Myxococcales - Bergey’s Manual of Systematic Bacteriology [22]. Morphology of fruiting bodies and colonies varies according to myxobacterial genera/species as follows [21]:

#### 2.3.2. 16S RNA analysis

##### DNA extraction

Myxobacterial was dispersed in 200 µL phosphate buffer. 20 µL Proteinase K and 200 µL cell lysis buffer were added and incubated at 72°C for 10 min. Continuously, 200 µL of absolute ethanol was added. The mixture was transferred on the silica column preset in 2-mL Eppendorf. After centrifugation at 13,000 rpm for 1 min, the liquid in the tube was removed, and the silica column contained DNA was put back in the tube. The DNA purification was repeated with 500 µL wash buffer 1 and 500 µL wash buffer 2 and centrifuged at 13,000 rpm for 1 min. The DNA in the column was resuspended with 50 µL elution buffer and incubated at room temperature for 1 min. After centrifuging for 1 min, the DNA solution was collected and used for PCR reaction [23].

##### The 16S rDNA amplification

16S rRNA gene was amplified using the 27F and 1492R primers. The final volume of PCR was 25 µl, including 5 µl of DNA, 1X reaction buffer (Tris KCl-MgCl_2_), 5 mmol MgCl_2_, 1 mmol dNTP, 0.4 µmol of each primer, and MyTaq HS DNA Polymerase (5 U/µL). The PCR program was initial denaturation at 95 °C for 5 min and 30 cycles (denaturation at 95 °C for 80 s, annealing at 57 °C for 30 s, and extending at 72 °C for 80 s) [24]. The PCR products were purified with DNA Genome Extraction Kit and sent for sequencing at Axil Scientific Pte Ltd, Singapore.

#### 2.3.3. Phylogenetic analysis

The sequences were overlapped using the Lasergene program. The phylogenetic neighbours were completed by the Basic Local Alignment Search Tool (Nucleotide BLAST) program and submitted to Gen Bank. The phylogenetic tree was constructed using the maximum likelihood method with 1000 replicates of bootstrap. All these programs were analyzed by MEGA software version 11 [24].

### 2.4. Preparation of myxobacterial crude extracts

The purified isolates were inoculated in 100-mL peptone media and shaken at 180 rpm at room temperature for ten days. 1-2% Amberlite XAD-16N adsorbent resin was added on the 4^th^-day of cultivation. After fermentation, the resin was collected by filtering, extracted with 80 mL acetone and then with 80 mL methanol, each for 2 hours. The organic solvents were evaporated in a vacuum device to a final volume of 1-mL crude extracts and stored at 4-8°C [25].

### 2.5. Evaluation of antioxidant activity

#### 2.5.1. DPPH assay

The DPPH (2,2-diphenyl-1-picryl-hydrazyl-hydrate, Sigma) stock solution 1 mmol was diluted with a solvent to achieve an absorbance of 0.680 (± 0.02) at 517 nm. The mixture was prepared by mixing 100 μL DPPH solution, 80 μL methanol, and 20 μL myxobacterial extract. The absorbance was recorded at a wavelength of 517 nm [26].

#### 2.5.2. ABTS radical cation scavenging activity

7 mmol ABTS (2,2’-azinobis-(3-ethylbenzothiazoline-6-sulfonic acid, Sigma) stock solution was mixed with 2.45 mmol potassium persulfate (K_2_S_2_O_8_) solution, which produced ABTS radical cation (ABTS^•+^). The final volume included 20 µL crude extract, 100 µL ABTS^•+^ solution (A_734 nm_ = 0.700 ± 0.020), and 80 µL the mixture of methanol and distilled water (1:1, v/v). The absorbance was measured at a wavelength of 734 nm. The well, containing 100 μL ABTS^•+^ solution, 50 μL methanol, and 50 μL distilled water, was negative control [27].

The IC_50_ values (the extract concentration that requires scavenging of 50% free radicals, µg/mL) were predicted from the regression model and used to express the antioxidative activities. Ascorbic acid and Trolox were used as the standards for establishing the ratio of IC_50 (sample)_ to IC_50 (standard)_ in DPPH and ABTS methods, respectively [28]. All measurements were carried out in triplicate and averaged.

### 2.6. Evaluation of antimicrobial activity

#### 2.6.1. Test microorganisms

Methicillin-susceptible *Staphylococcus aureus* ATCC 25923, Methicillin-resistant *Staphylococcus aureus* ATCC 43300 (MR), *Streptococcus faecalis* ATCC 29212 (Sf), *Escherichia coli* ATCC 25922 (Ec), *Pseudomonas aeruginosa* ATCC 27853 (Pa), *Candida albicans* ATCC 10231 (Ca), *Aspergillus niger* ATCC 16404 (An); Penicillium sp. (Pe), Mucor sp. (Mu), and Rhizopus sp. (Rh) were supplied by University of Medicine and Pharmacy at Ho Chi Minh City.

#### 2.6.2. Microdilution method

Muller Hinton broth and RPMI-1640 medium were respectively used for bacterial and filamentous fungi strains, and the supplement of 2% glucose was used for yeast *C. albicans*. Minimum inhibitory concentrations (MICs) of extracts were determined by the microdilution method using 96-well microtiter plates. The extracts were diluted in DMSO with the highest concentration in well 1 was 512 µg/mL, followed by dilution of ½ to obtain a decreasing range of concentrations. Add 50 µL test microorganism inocula; and 30 µL resazurin at 0.015 mg/mL as an indicator for cell viability; plates were incubated at 35°C for 16-48 hours [29]. The MIC endpoints were observed visually when viable cells changed their colour from purple to pink [30].

## 3. Results

### 3.1. Isolation and identification

Soil samples were mainly loam, peat, and silt soils that were collected in rice fields or near the root of large trees in fruit gardens in Vietnam. All of the strains isolated from soil were inoculated on VY2 medium, and then the morphology of fruiting bodies, myxospores, and vegetative cells was observed under a stereoscopic microscope. In which, based on morphological characteristics and 16S ribosomal RNA gene sequence, 43 strains were determined entirely to belong to Myxococcales, including 15 (34.9%), 16 (37.2%), and 12 (27.9%) strains from *E. coli* bait/WCX method, rabbit dung, and filter paper methods, respectively (Figure 1).

**Figure 1.**
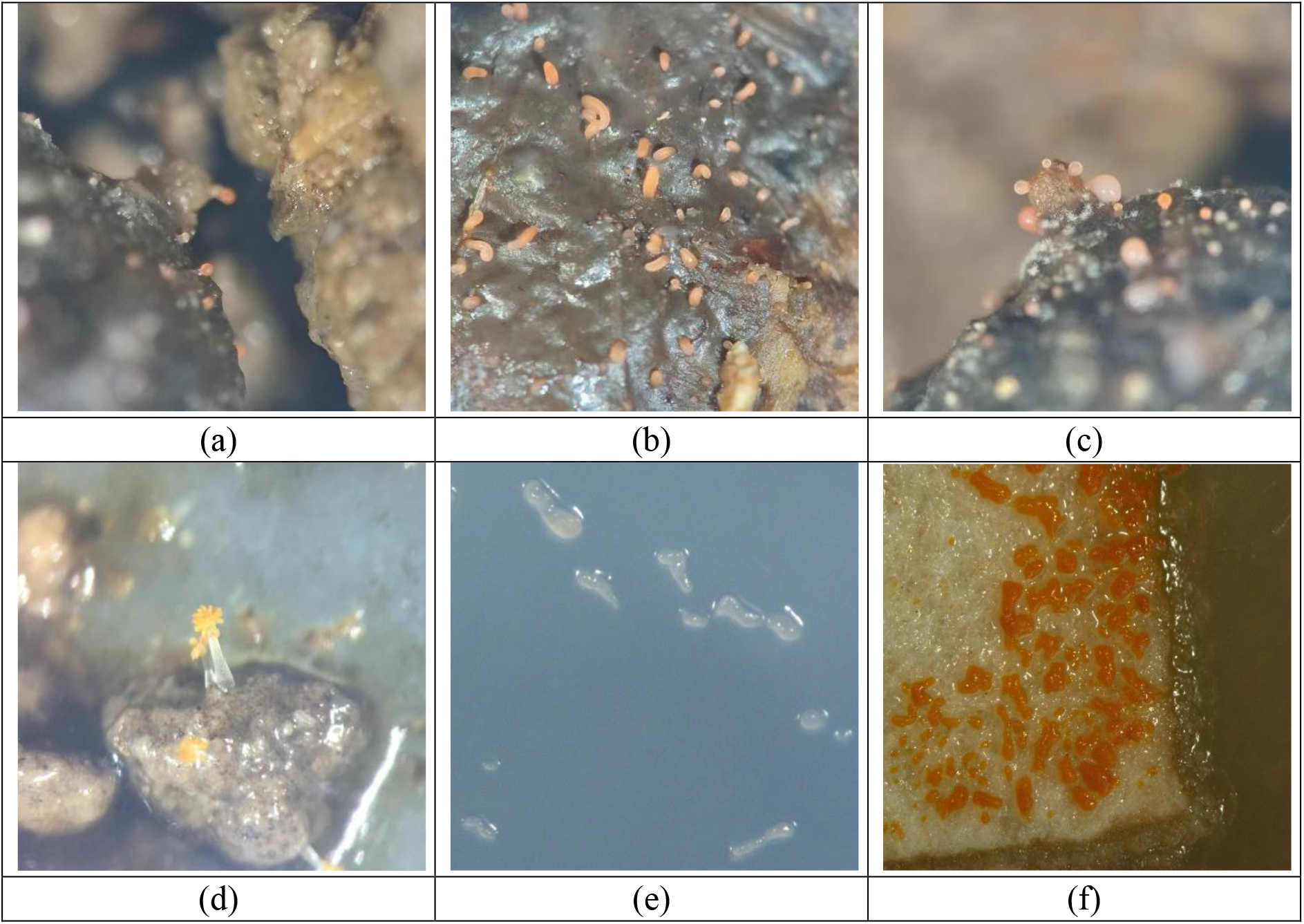
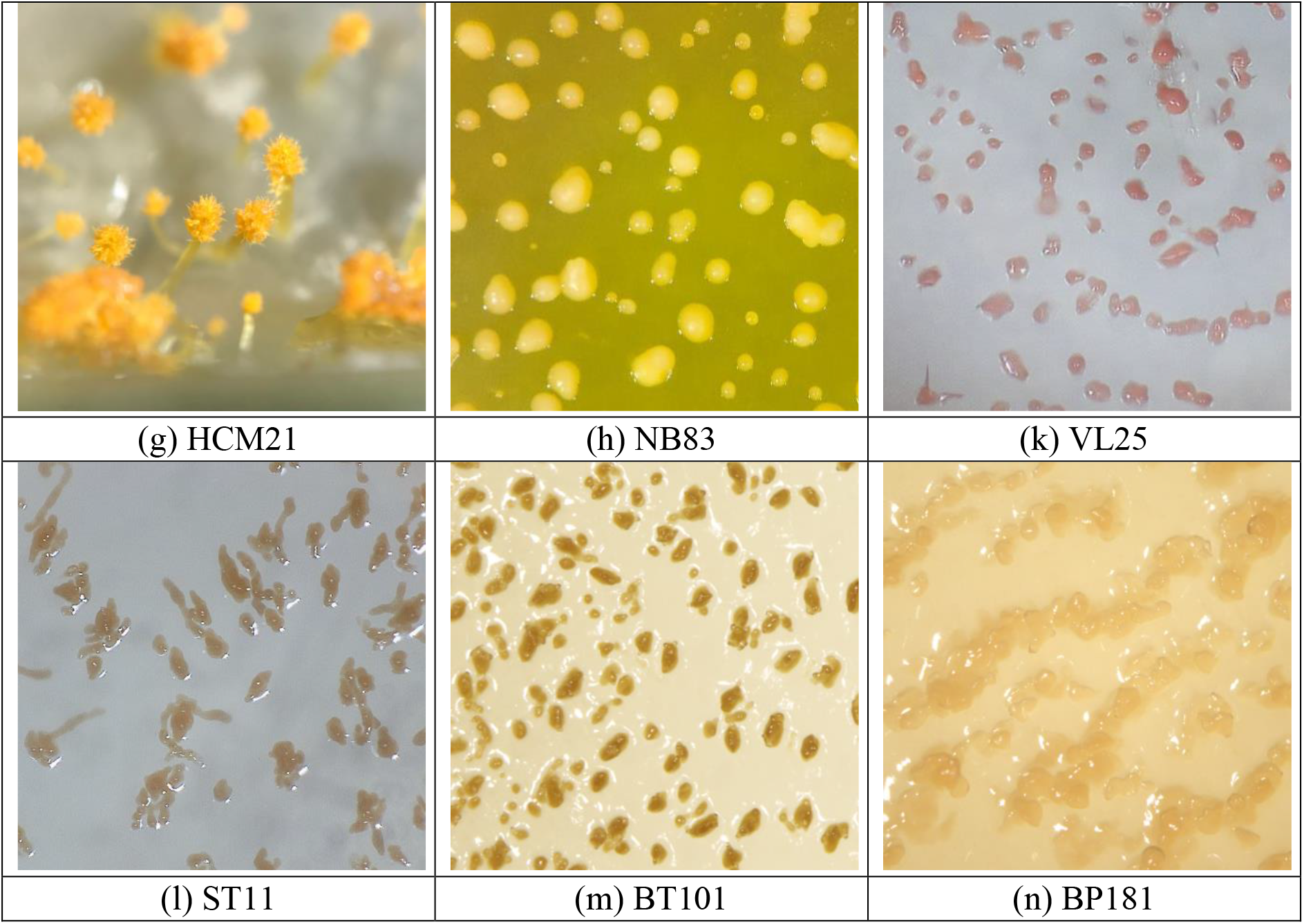
Myxobacterial fruiting bodies Fruiting bodies on rabbit dung (a-c), on WCX agar (d, e), and on filter paper (f); Fruiting bodies of isolates on VY2 medium (g-n)

The morphology of the colonial edge, the opaque and the radial pattern; the colour and shape of fruiting bodies, aggregate or solitary structure; and the size of vegetative cells were observed on the dissecting and optical microscopes. Vegetative cells were generally characterized by slender rods and rounded ends, approximately 3-12 µm in length (Figure 2).

**Figure 2.**
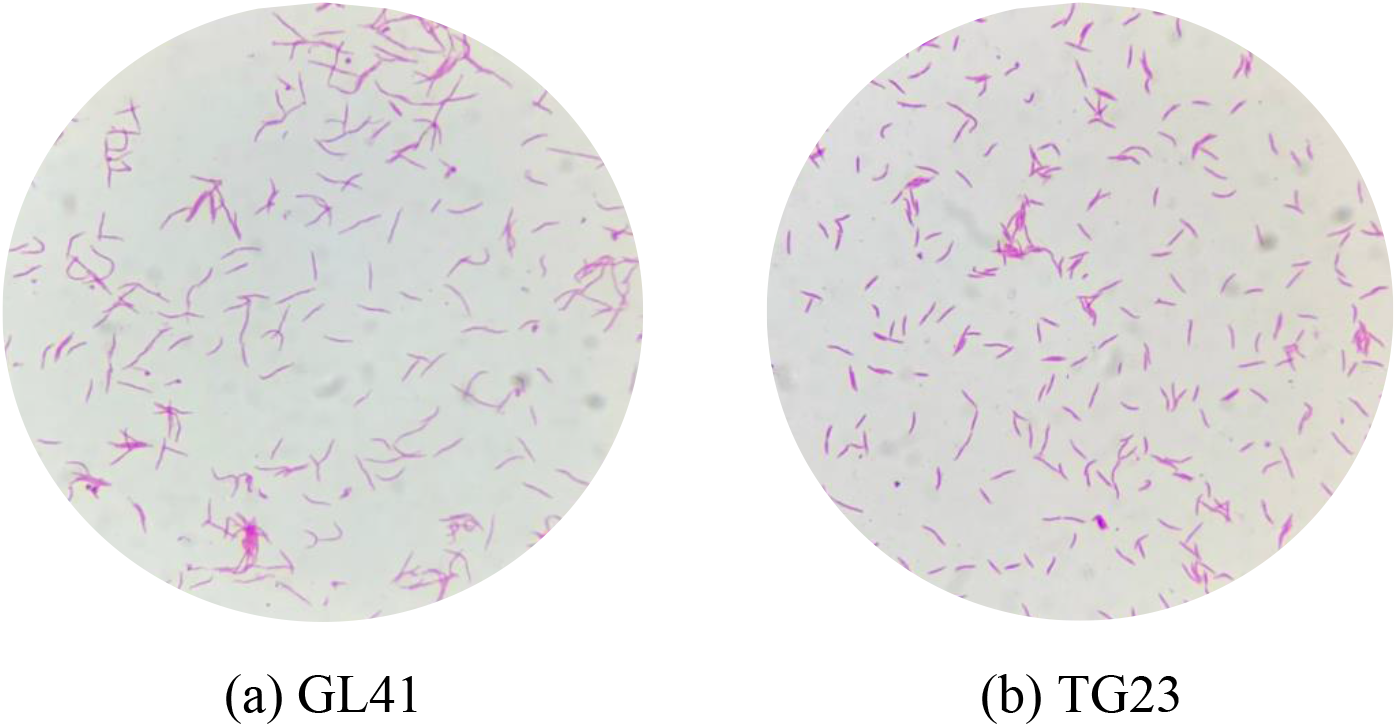
Morphology of Gram-negative Myxobacteria from GL41 (a) and TG23 (b)

Continuously, 16S RNA gene sequences were analyzed. Forty-three strains were designated into seven genera: *Angiococcus* (1), *Archangium* (3), *Chondromyces* (2), *Corallococcus* (15), *Cystobacter* (1), *Melittangium* (1), and *Myxococcus* (20) (Figure 3 and Table 3).

**Figure 3.**
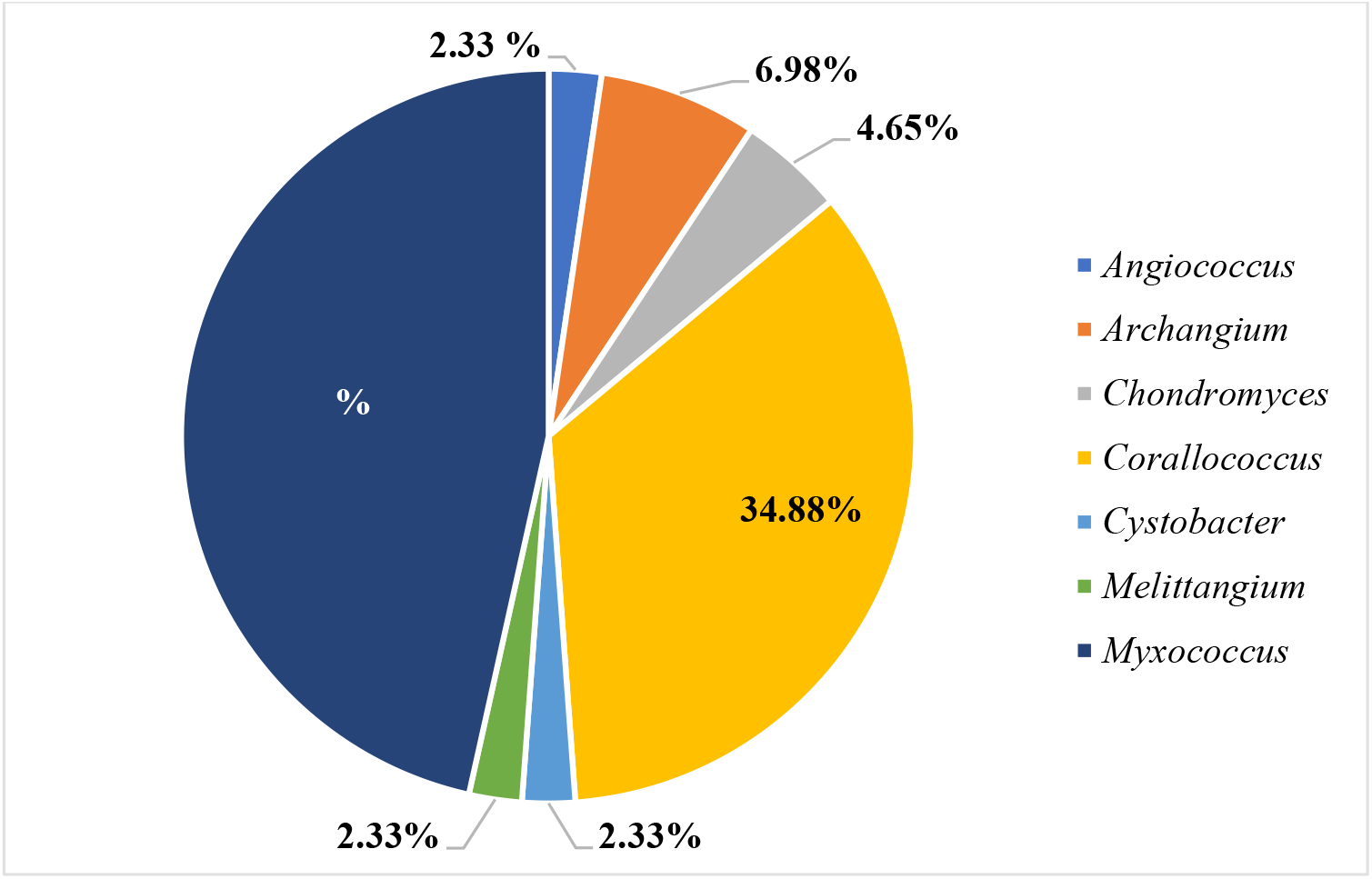
The number of myxobacterial strains

**Table 3.**
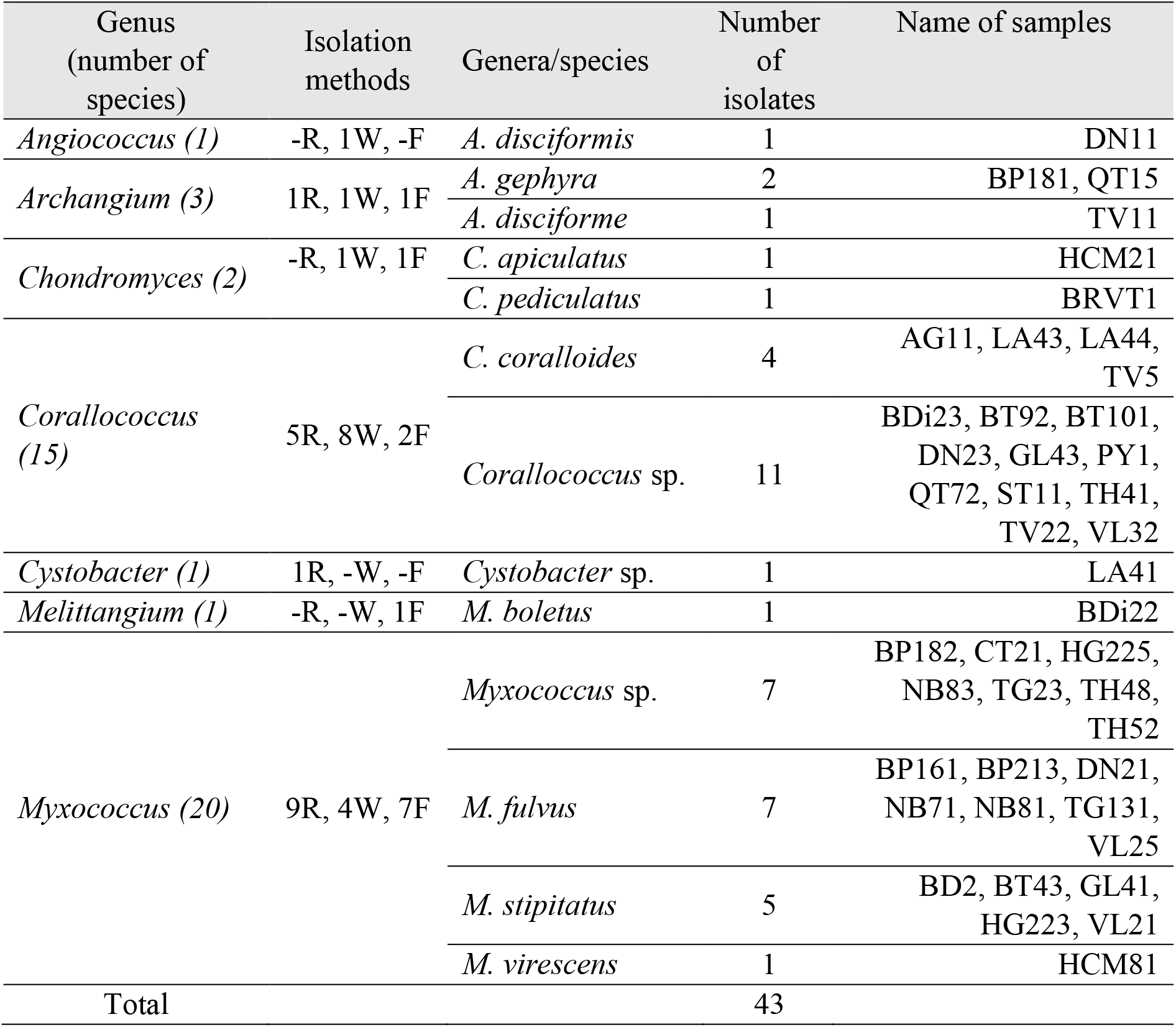
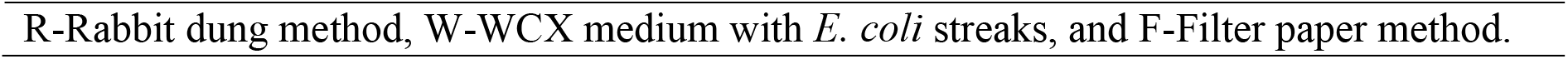
Identification of isolates according to 16S rDNA analysis

Forty-three strains show a high similarity (99.65-100%) with the above seven genera. The phylogenetic tree was built to determine the genetic relationships between isolates and the known strains (Figure 4).

**Figure 4.**
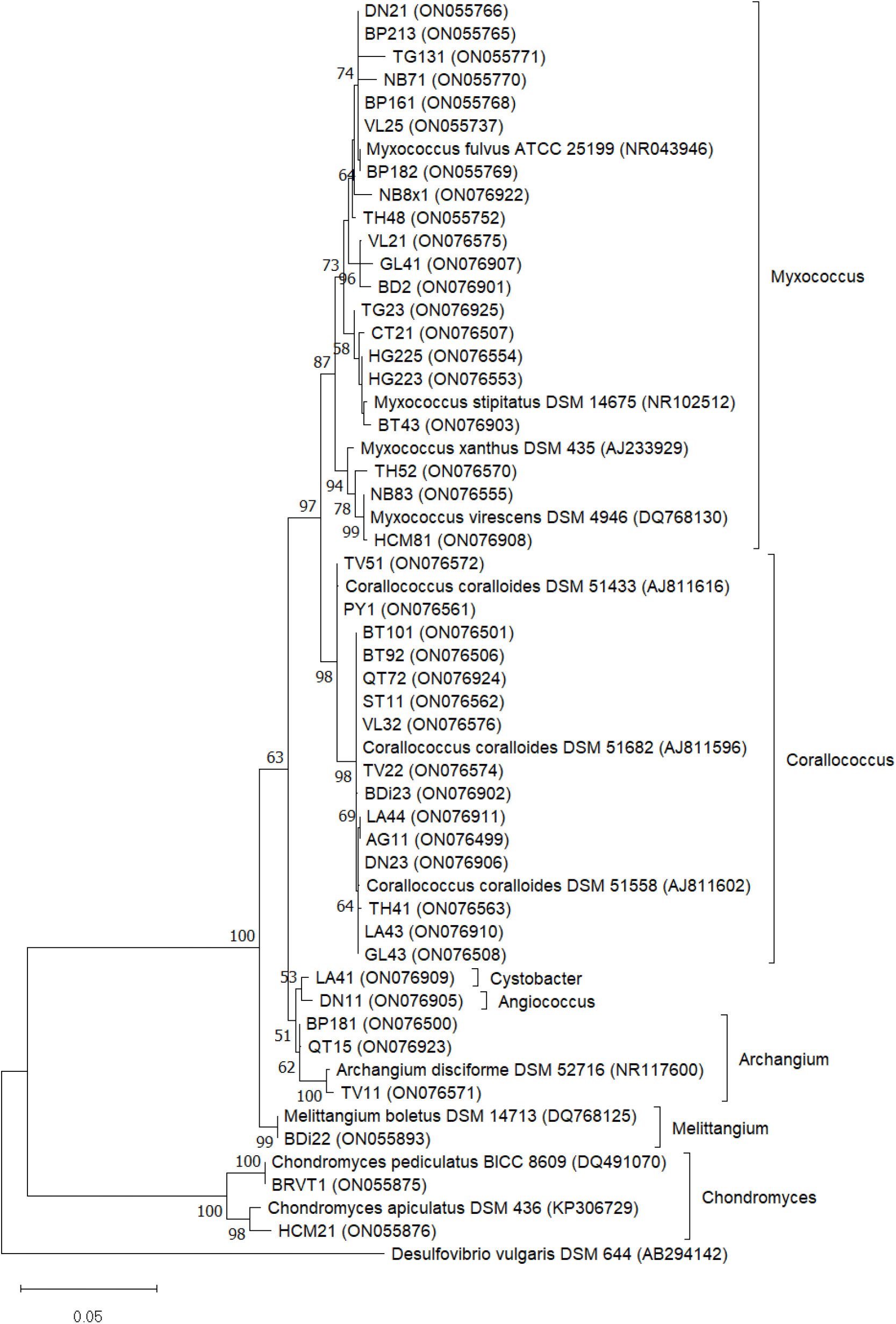
Phylogenetic tree of 43 strains based on 16S rDNA gene sequences

Consequently, based on the morphology and the 16S rRNA gene sequences, 43 strains were classified into seven genera, three families (Myxococcaceae, Archangiaceae, and Polyangiaceae), and two suborders (Cystobacterineae and Sorangiineae). Among the 43 strains of myxobacteria, 24 strains were identified to the species level, while 19 strains were assigned to the genus level (Table 3).

### 3.2. Screening antioxidant activity

Data on the 43 myxobacterial extract concentrations that neutralize 50% DPPH^•^ ranged from 27.39 ± 1.74 to 249,43 ± 6,17 µg/mL. The strain with the highest activity is TH41 (IC_50_ = 27.39 ± 1.74 µg/mL, ratio = 2.51), followed by TG131, VL32, CT21, and GL41 with the 50% scavenger concentration of 40.28 ± 1.13 μg/mL (ratio = 3.70), 50.87 ± 1.33 μg/mL (ratio = 4.67), 52.34 ± 1.47 μg/mL (ratio = 4.80), and 57.24 ± 1.52 µg/mL (ratio = 5.25), respectively (Table 4).

**Table 4.**
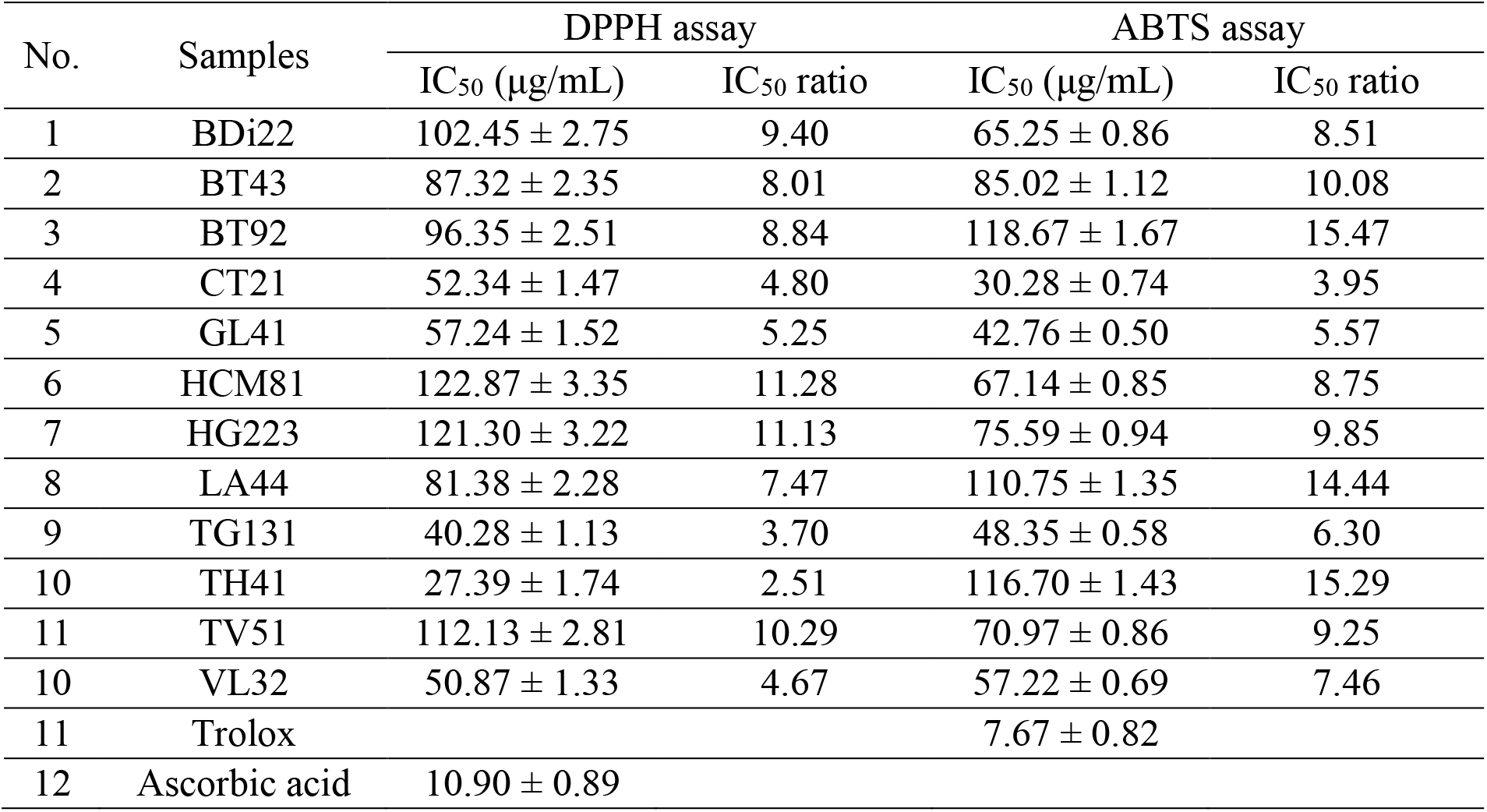
High antioxidant activities of some myxobacterial strains

| No. | Samples | DPPH assay |  | ABTS assay |  |
| --- | --- | --- | --- | --- | --- |
| | | $\text{IC}_{50}$ ($\mu\text{g/mL}$) | $\text{IC}_{50}$ ratio | $\text{IC}_{50}$ ($\mu\text{g/mL}$) | $\text{IC}_{50}$ ratio |
| 1 | BDi22 | $102.45 \pm 2.75$ | 9.40 | $65.25 \pm 0.86$ | 8.51 |
| 2 | BT43 | $87.32 \pm 2.35$ | 8.01 | $85.02 \pm 1.12$ | 10.08 |
| 3 | BT92 | $96.35 \pm 2.51$ | 8.84 | $118.67 \pm 1.67$ | 15.47 |
| 4 | CT21 | $52.34 \pm 1.47$ | 4.80 | $30.28 \pm 0.74$ | 3.95 |
| 5 | GL41 | $57.24 \pm 1.52$ | 5.25 | $42.76 \pm 0.50$ | 5.57 |
| 6 | HCM81 | $122.87 \pm 3.35$ | 11.28 | $67.14 \pm 0.85$ | 8.75 |
| 7 | HG223 | $121.30 \pm 3.22$ | 11.13 | $75.59 \pm 0.94$ | 9.85 |
| 8 | LA44 | $81.38 \pm 2.28$ | 7.47 | $110.75 \pm 1.35$ | 14.44 |
| 9 | TG131 | $40.28 \pm 1.13$ | 3.70 | $48.35 \pm 0.58$ | 6.30 |
| 10 | TH41 | $27.39 \pm 1.74$ | 2.51 | $116.70 \pm 1.43$ | 15.29 |
| 11 | TV51 | $112.13 \pm 2.81$ | 10.29 | $70.97 \pm 0.86$ | 9.25 |
| 10 | VL32 | $50.87 \pm 1.33$ | 4.67 | $57.22 \pm 0.69$ | 7.46 |
| 11 | Trolox | | | $7.67 \pm 0.82$ | |
| 12 | Ascorbic acid | $10.90 \pm 0.89$ | | | |

All samples showed a high capacity to scavenge ABTS^•+^ cation radicals. The IC_50_ values from 43 extracts significantly differed between myxobacterial extracts, varying from 30.28 ± 0.74 to 197,36 ± 2,22 µg/mL. The CT21 extract exhibits the highest activity (equivalent 30.28 ± 0.74 µg/mL, ratio = 3.95), followed by GL41, TG131, VL32, and BDi22 extracts for the values of 42.76 ± 0.50 μg/mL (ratio = 5.57), 48.35 ± 0.58 μg/mL (ratio = 6.30), 57.22 ± 0.69 μg/mL (ratio = 7.46), and 65.25 ± 0.86 μg/mL (ratio = 8.51), respectively (Table 4).

In detail, the results of antioxidative evaluation exhibited that crude extract from CT21 was the highest antioxidant activity in both assays (IC_50_ = 52.34 ± 1.47 µg/mL, ratio = 4.80 for DPPH assay, and 30.28 ± 0.74 µg/mL, ratio = 3.95 for ABTS assay), proving that the scavenging capacity of antioxidants towards DPPH^•^ and ABTS^+•^. The other potential strains were TG131 that the measured IC_50_ values were 40.28 ± 1.13 µg/mL (ratio = 3.70) and 48.35 ± 0.58 µg/mL (ratio = 6.30) for DPPH, ABTS assays, respectively. The GL41 also demonstrated the low half-maximal inhibitory concentration corresponding to 57.24 ± 1.52 µg/mL (ratio = 5.25) in DPPH and 42.76 ± 0.50 µg/mL (ratio = 5.57) in ABTS.

### 3.3. Evaluation of antimicrobial activities

The results of MIC of myxobacterial extracts showed that 100% extracts on P-medium against at least one of ten test microorganisms with broadly varied concentrations ranging from 1 to 512 µg/mL.

In this study, 40, 23, and 18 strains produced bioactive metabolites against MRSA, MSSA and *S. faecalis*, respectively. However, there are only 16 strains against *P. aeruginosa*, and only one strain was found to be sensitive to *E. coli*. All strains show antifungal effects, with 55.8 - 86.0% of the isolates inhibiting yeast and moulds, in which the highest ratio (86% of extracts) showed activity against Mucor sp., and the lowest was 55.8% against *C. albicans*. The susceptibility of extracts on Gram-positive bacteria and mould generally is better than on Gram-negative bacteria and yeast, respectively (Table 5).

**Table 5.** Antimicrobial activity of isolated genera (µg/mL)

| Genera | Test microorganisms |  |  |  |  |  |  |  |  |  |
| --- | --- | --- | --- | --- | --- | --- | --- | --- | --- | --- |
|  | MR | MS | Sf | Ec | Pa | Ca | Mu | Rh | Pe | An |
| *Angiococcus* (n = 1) | 1 | - | - | - | - | - | 1 | 1 | - | - |
| *Archangium* (n = 3) | 3 | 3 | 2 | - | 1 | 2 | 2 | 2 | - | 1 |
| *Chondromyces* (n = 2) | 1 | 1 | 1 | - | 1 | - | 1 | 2 | 2 | 2 |
| *Corallococcus* (n = 15) | 15 | 7 | 4 | - | 5 | 7 | 10 | 13 | 12 | 15 |
| *Cystobacter* (n = 1) | 1 | - | - | - | - | - | - | - | 1 | 1 |
| *Melittangium* (n = 1) | 1 | 1 | - | - | - | 1 | - | 1 | 1 | 1 |
| *Myxococcus* (n = 20) | 18 | 11 | 11 | 1 | 9 | 14 | 17 | 18 | 14 | 16 |
| Total (n = 43) | 40 | 23 | 18 | 1 | 16 | 24 | 31 | 37 | 30 | 36 |
| (%) | 93.0 | 53.5 | 41.9 | 2.3 | 37.2 | 55.8 | 72.1 | 86.0 | 69.8 | 83.7 |
| MR: MRSA, MS: MSSA, Sf: *S. faecalis*, Ec: *E. coli*, Pa: *P. aeruginosa*, Ca: *C. albicans*, Mu: *Mucor* sp., Rh: *Rhizopus* sp., Pe: *Penicillium* sp., and An: *A. niger* |  |  |  |  |  |  |  |  |  |  |

Significantly active strains (6/9 strains) belonged to the Myxococcus genus. Strain GL41 was found to be the highest inhibitory effect on MRSA, MSSA, *S. faecalis, C. albicans*, and *A. niger* with MIC were 1 µg/mL. All of the test microorganisms were indicated to be susceptible to GL41. This is also the only extract that sensitive to *E. coli* (Table 6).

**Table 6.** MIC of isolates against test microorganisms (μg/mL)

| Isolates | MR | MS | Sf | Ec | Pa | Ca | Rh | Mu | Pe | An |
| --- | --- | --- | --- | --- | --- | --- | --- | --- | --- | --- |
| BD2 | 4 | 4 | 32 | - | 512 | 8 | 4 | 8 | 2 | 1 |
| BDi22 | 128 | 512 | - | - | - | 16 | 32 | 64 | 1 | 1 |
| BP161 | 4 | 1 | 16 | - | 512 | 128 | 64 | 128 | - | - |
| BP181 | 1 | 1 | 4 | - | 512 | 64 | 128 | 256 | - | - |
| CT21 | 128 | 64 | 512 | - | - | 8 | 16 | - | 2 | 1 |
| GL41 | 1 | 1 | 1 | 64 | 128 | 1 | 16 | 16 | 8 | 1 |
| HG225 | 16 | 64 | 64 | - | 512 | 32 | 16 | 16 | 1 | 1 |
| NB71 | 4 | 1 | 16 | - | 512 | 16 | 32 | 16 | 1 | 1 |
| VL32 | 64 | 256 | - | - | - | 512 | 128 | 64 | 1 | 4 |
| Note: MR: MRSA, MS: MSSA, Sf: *S. faecalis*, Ec: *E. coli*, Pa: *P. aeruginosa*, Ca: *C. albicans*, Mu: *Mucor* sp., Rh: *Rhizopus* sp., Pe: *Penicillium* sp., and An: *A. niger* |  |  |  |  |  |  |  |  |  |  |

## 4. Discussion

Many scientific reports over the centuries have proven the enormous source of myxobacterial-derived metabolites. Affected by the tropical climatic zone and geographic biodiversity, Vietnam could be predicted the microbial potential of Myxococcales. The current approach is the first publication that studies the isolation, identification, and bioactive screening of myxobacteria in Vietnam.

### 4.1. Myxobacterial isolation, identification and phylogenetic analysis

The results showed that forty-three myxobacterial strains were isolated and identified from 20 provinces and cities in Vietnam. These isolates were identified by morphology and 16S rRNA sequence. The rabbit dung method showed the highest yield of myxobacterial isolation, which accounted for 37.2%. The *E. coli* baiting method was 34.9%, and the lowest number was 27.9% representing the selective isolation of cellulose-degradable strains.

Rabbit dung technique with high isolation efficiency and easy purification is a remarkable difference compared to previous research. In detail, the ratio of purified strains from rabbit dung (37.2%) was much higher than 3.4% in saline-alkaline soils in Xinjiang, China (2013) [31], but lower than 75.0% in Israel study (2005) [32], partly due to the isolation has been affected by different soil physical properties. Using a large amount of soil in the rabbit dung method allowed to improve the isolating probability. The myxobacteria could induce naturally organic substrates and glide toward the rabbit dung to form fruiting bodies on the autoclaved pellets. Likewise, on WCX agar, myxobacteria also perform the gliding movement from the soil sample along with the *E. coli* smears and easily recognize it. However, some other contaminants, such as alphaproteobacteria or deltaproteobacteria, can also create similar and confusing structures. In the the filter paper method, it requires a long time (2-4 weeks) for cellulose decomposition that is sufficiently visible to the naked eye, facilitating the overgrowth of infectious fungi and insects that interfere with further purification. Therefore, about 27.9% of isolates were obtained from pick-up fruiting bodies on dung pellets, but substantially higher than 2.9% of isolates in F. Gaspari et al. [32].

Except for some strains characterized by particular fruiting body structures (for example, Chondromyces sp.), it usually becomes difficult to distinguish the majority of myxobacteria from genus and species because the fruit body morphology is similar and the colour is often diverse, even though they change rapidly in laboratory conditions [23]. From that, the analysis of 16S gene sequence similarity can be effective. However, the data is not specific enough to determine the species level [31]. There may even be a certain similarity between closely related genera. In this study, molecular analysis showed that forty-three 16S rDNA sequences were assigned to seven genera, mainly belonging to the family Myxoccocaceae (Myxococcus, Corallococcus), suborder Cystobacterineae. This data is suitable with reports of Xianjiao Zhang et al. (2013) [31] and Siti Meliah et al. (2018) [23] with 43/58 and 7/10 strains belonging to Myxococcaceae, respectively.

### 4.2. Antioxidant activities

Although myxobacterial strains were isolated from different sources of soil properties or areas, the findings showed that approximately 100% of screened isolates exhibited high antioxidant activity with a considerable variation. The CT21 and GL41 extracts possess a potent DPPH^•^ free radical scavenger and indicate an excellent correlation to ABTS method. Compared with Trolox/ ascorbic acid, ratio values of IC_50, sample_: IC_50, reference_ changes from 3.95 - 5.57, suggested both crude extracts of CT21 and GL41 express exceptional antioxidant capacity.

The free radical scavenging properties of myxobacterial extracts are mainly due to the presence of endogenous compounds containing phenolic hydroxyl groups that have been known for the power neutralizing or reducing various oxidant agents. Myxobacteria generates phenolics as secondary metabolites that protect themselves from the damage induced by reactive oxygen species (ROS). As a part, these substances are also responsible for a broad collection of cytotoxic and antioxidant activities. Indeed, Mona Dehhaghi et al. (2018) summarized that myxobacterial extracts could decrease the formation of endogenous oxidative species and cell death [33].

### 4.3. Antimicrobial activities

Besides micro predator characteristics, Myxobacteria also synthesizes epibiotic compounds to compete with other natural microbiomes. All strains exhibited growth inhibition on at least one of ten microorganisms that inclined onto Gram-positive rather than Gram-negative strains, suitable with Ivana Charousová’s report (2017) [25]. The majority of the Myxococcus members showed considerably antimicrobial activity. Myxococcus genus was responsible for about 2/3 of the highest active strains, which could be elucidated by the most frequently isolated capacity from the natural substrate. In the report of F. Gaspari (2005), Myxococcus accounted for 62/97 isolated strains and 52/62 active strains [32]. Two of the best antimicrobial strains in a report by Ivana Charousová et al. exhibited a similarity of 99% with *M. xanthus* sequence [25].

The antifungal effect was generally better than bacteria. Juana Diez et al. (2012) supposed that about 54 and 29% of the biological compounds were antifungal or antibacterial agents, respectively [34]. Strain GL41 (identified as *Myxococcus stipitatus*) showed impressive activities that inhibit at the MIC values of 1 µg/mL on 5 bacterial and fungal strains. This strain also is one of three strains that possess the highest antioxidant properties. Objectively, some isolates were observed with insignificant effect, which may be explained by (i) one culture medium (P-medium) is not entirely optimized for producing the active secondary compounds. Therefore, bioactive screening of the extracts from many different media facilitates the appropriate evaluation, and (ii) each strain may possess other potential (anticarcinogenic) than oxidant and microbial inhibitory activities.

## 5. Conclusion

The study has described the isolation and phylogenetic relationship of myxobacteria in Vietnam. Besides, total extracts’ antioxidant and antimicrobial activities obtained from P-medium fermentation were determined. Further studies will be developed to evaluate the *in vitro* antitumor activities and fractional extraction of tremendous strains to identify the components responsible for bioactivity.

## Supporting information

https://mail.google.com/mail/u/0/#inbox

---

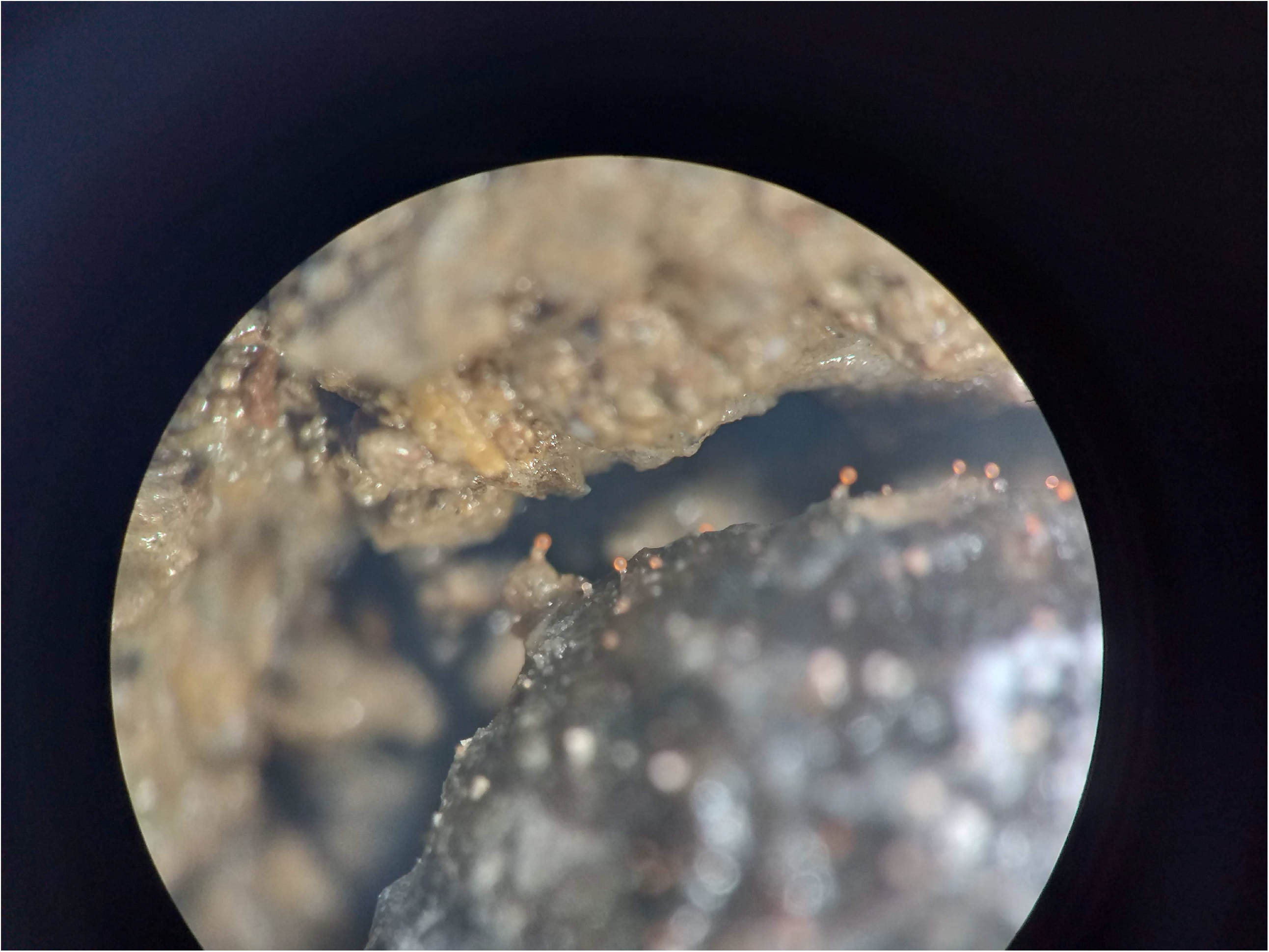

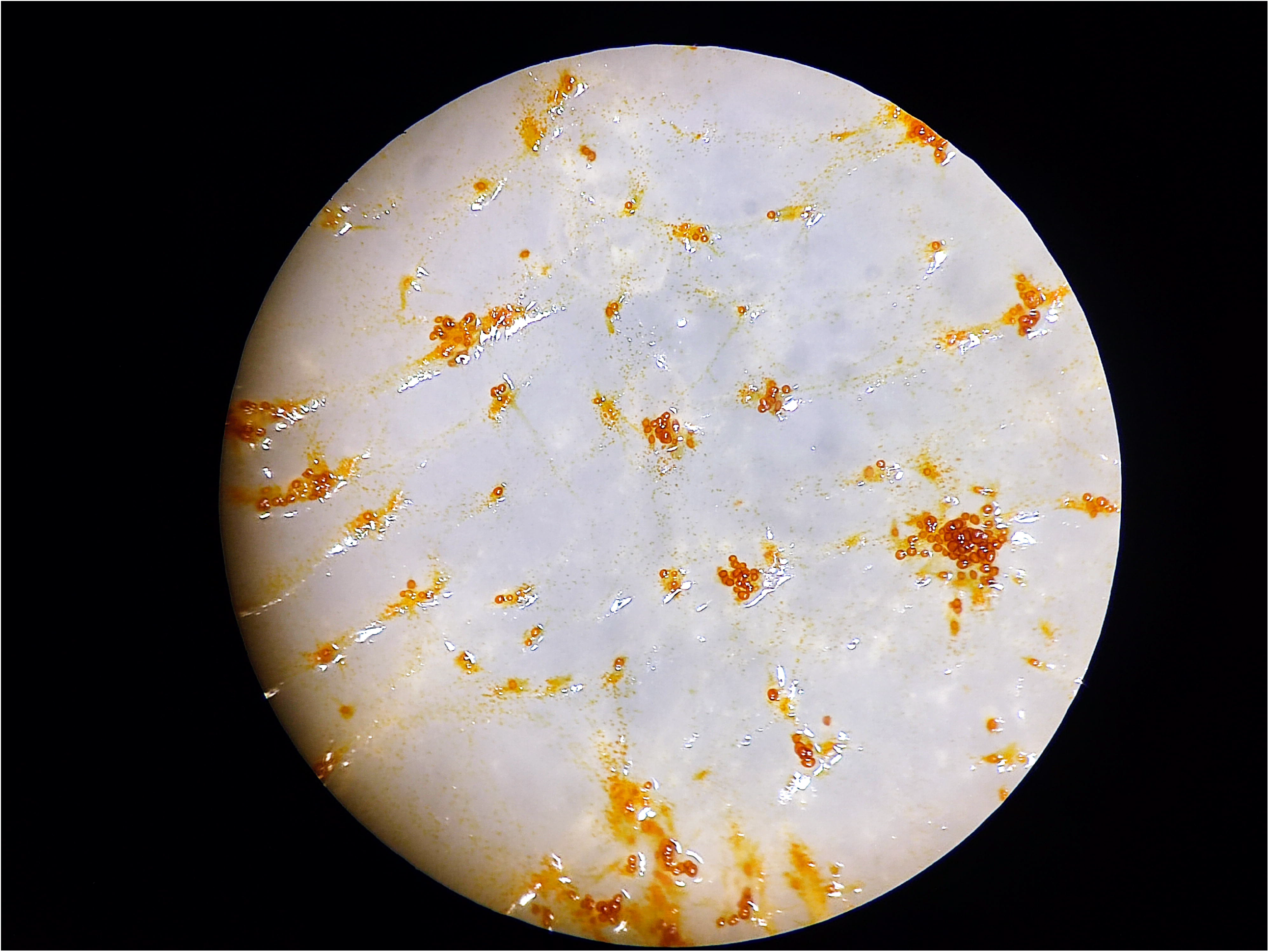

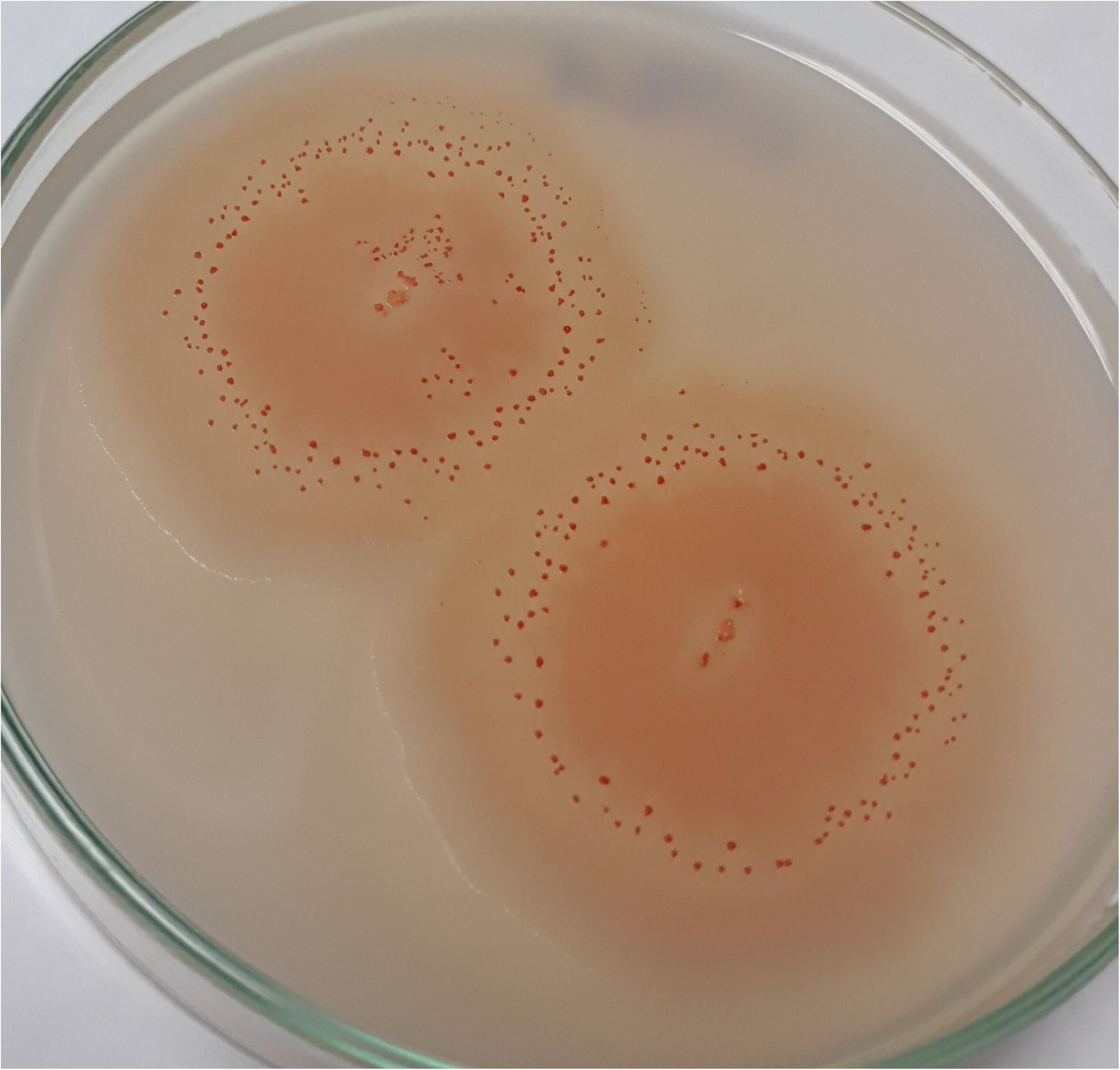

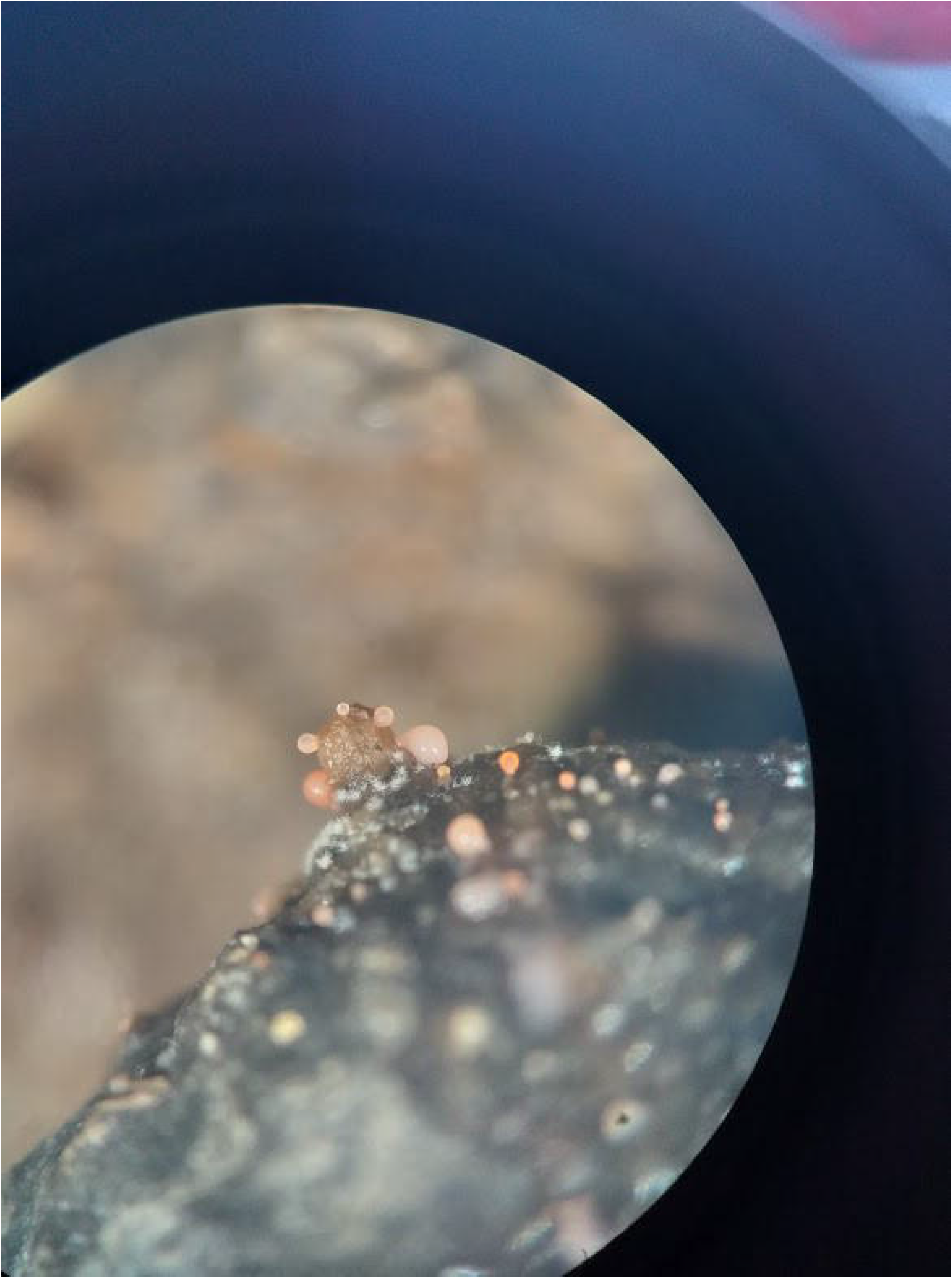

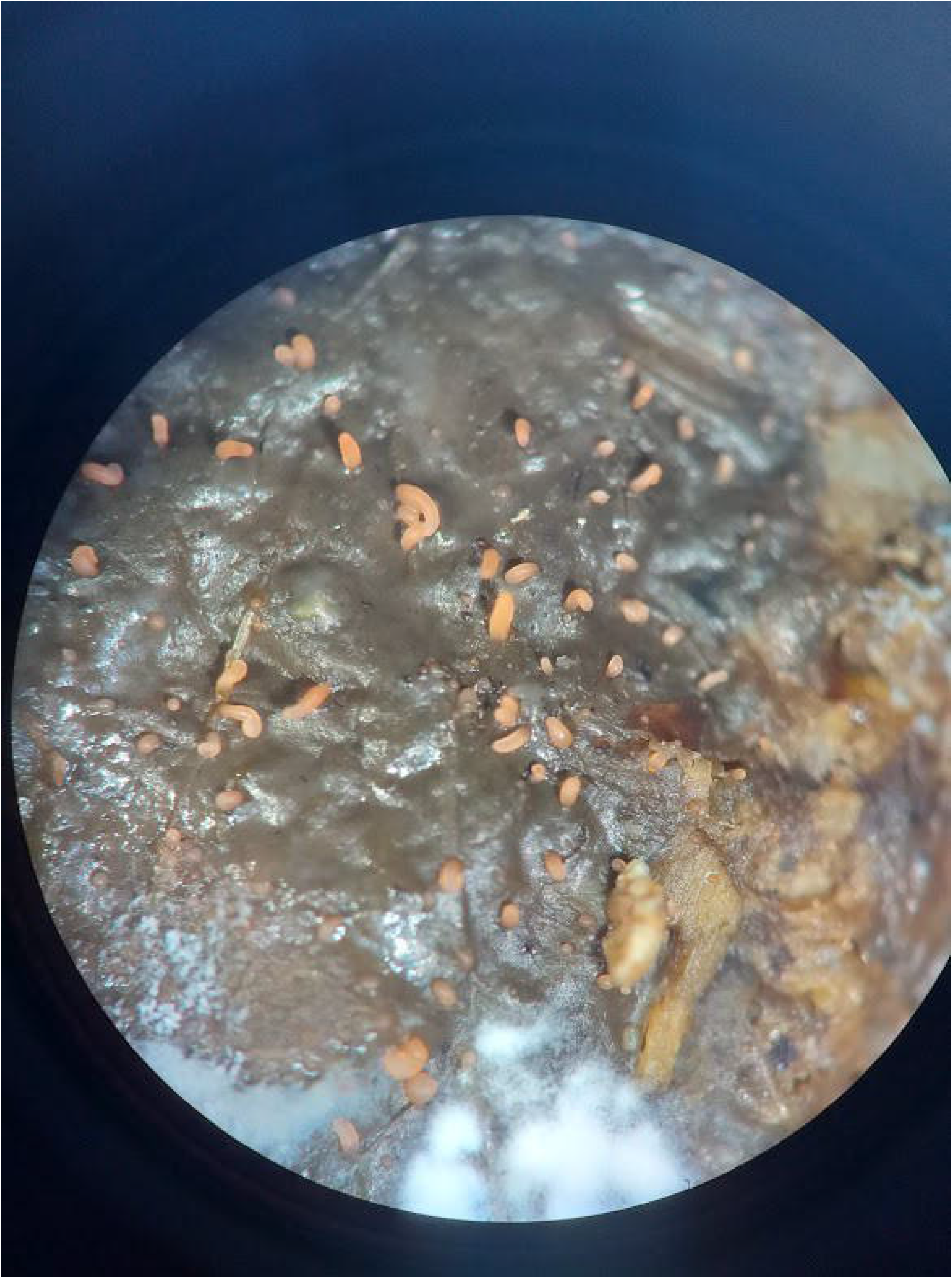

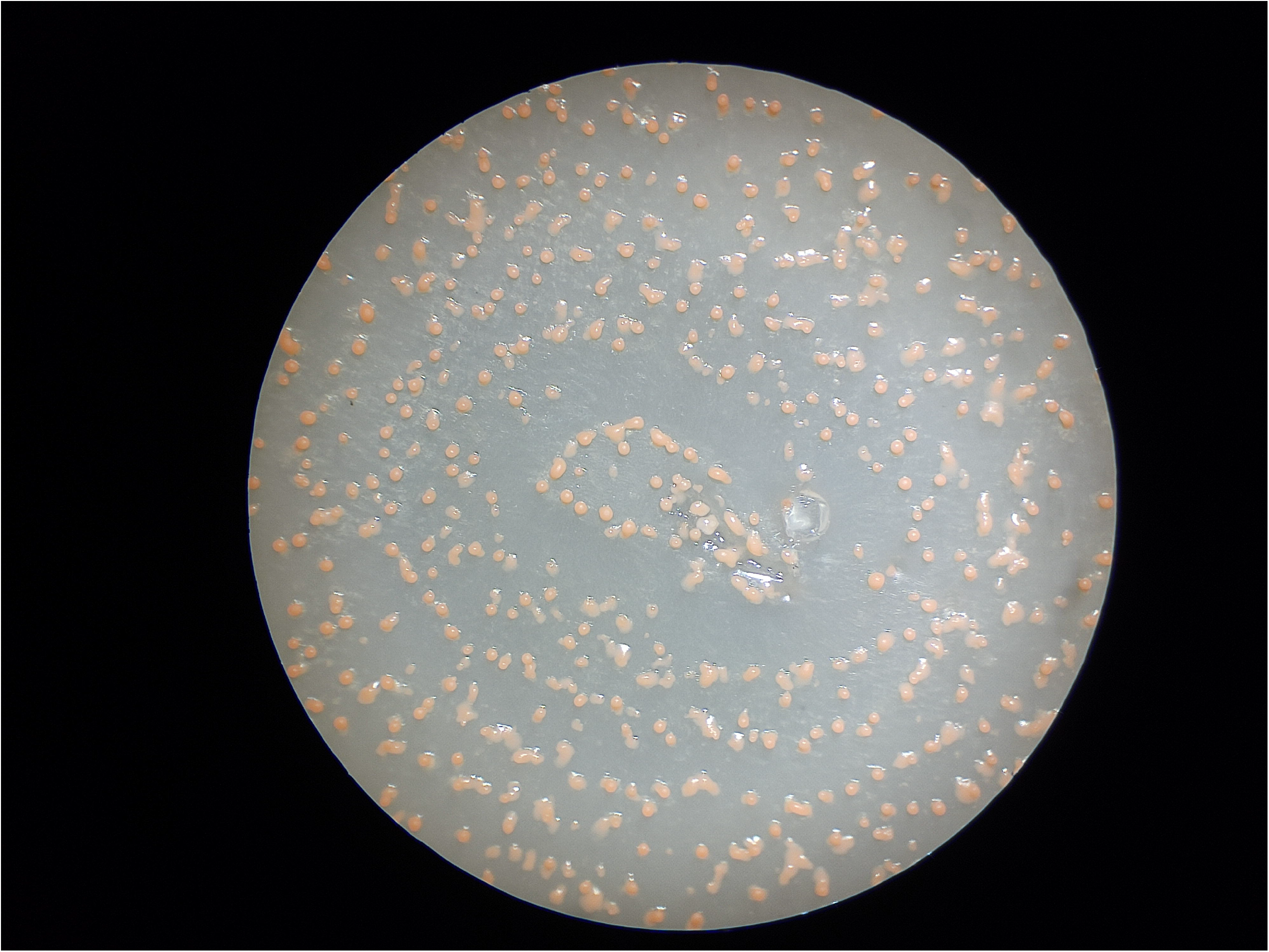

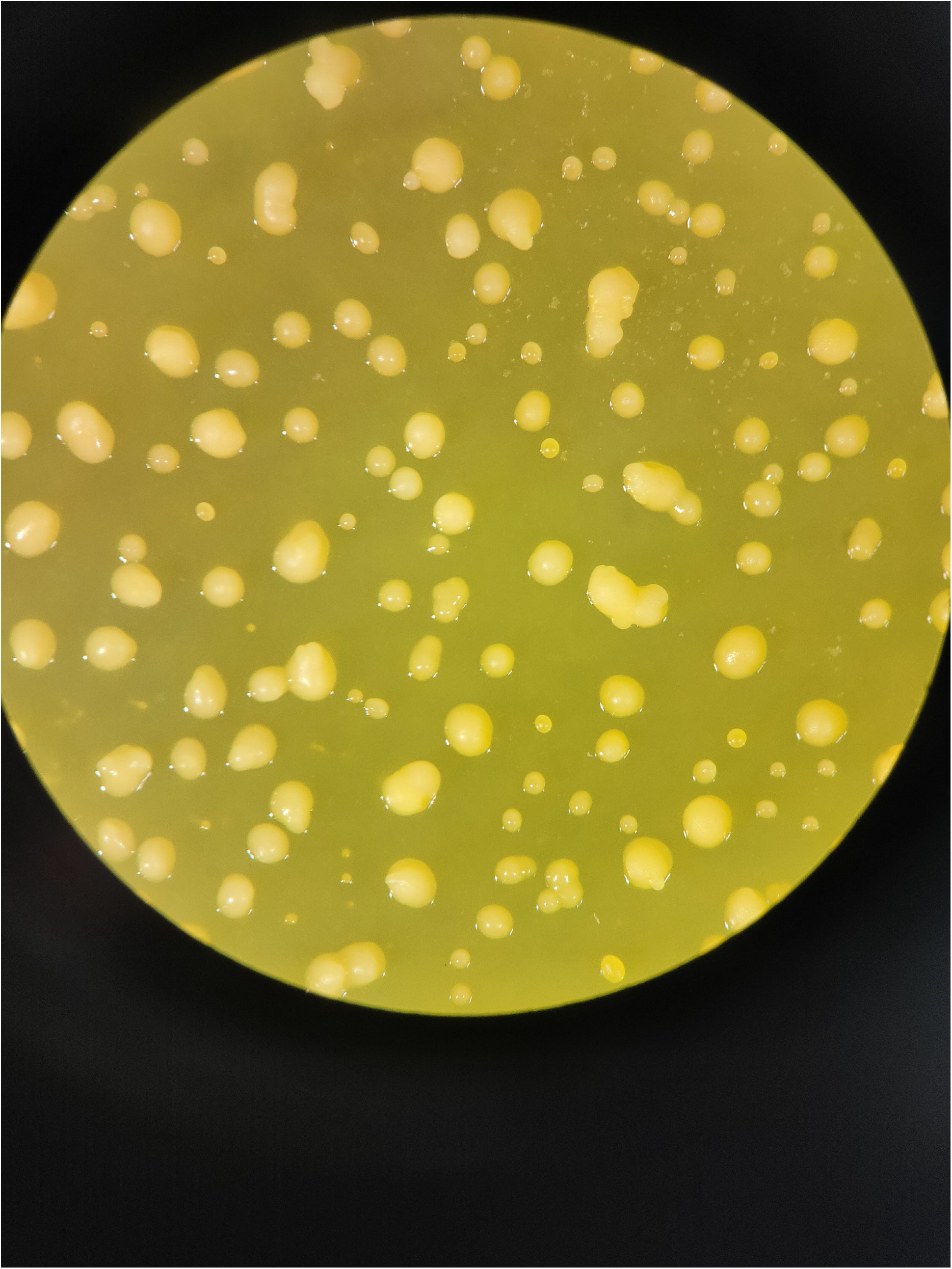

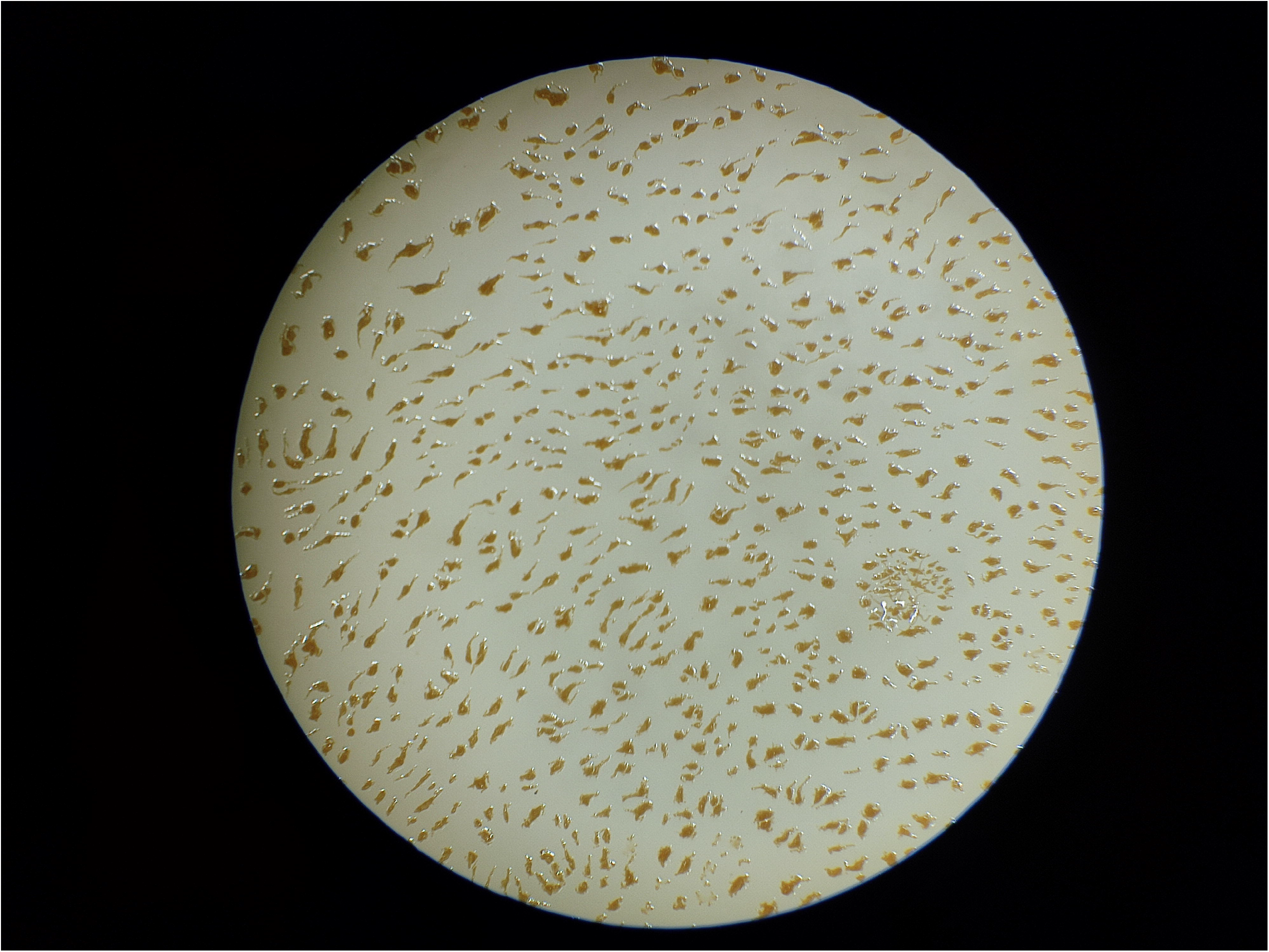

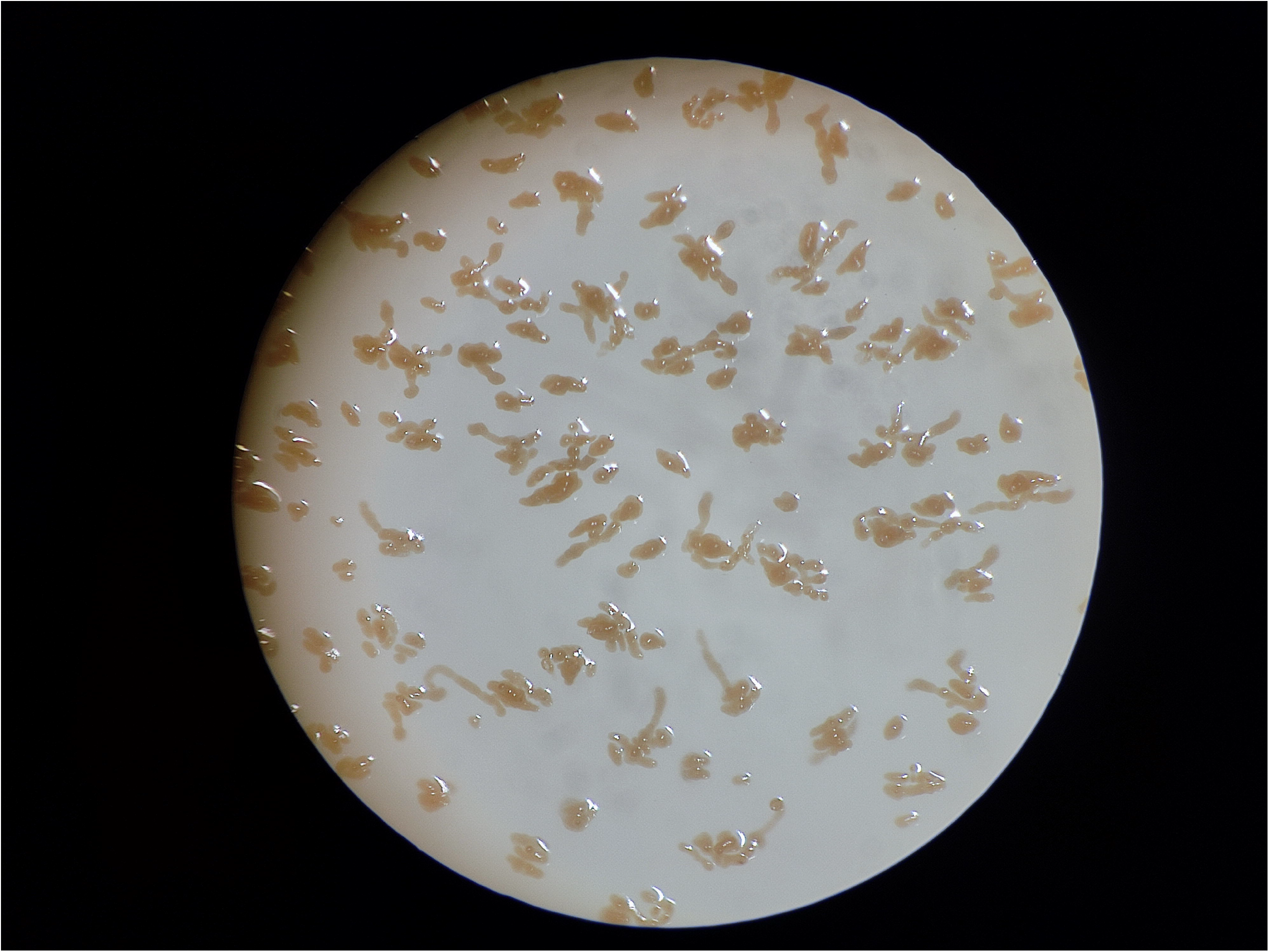

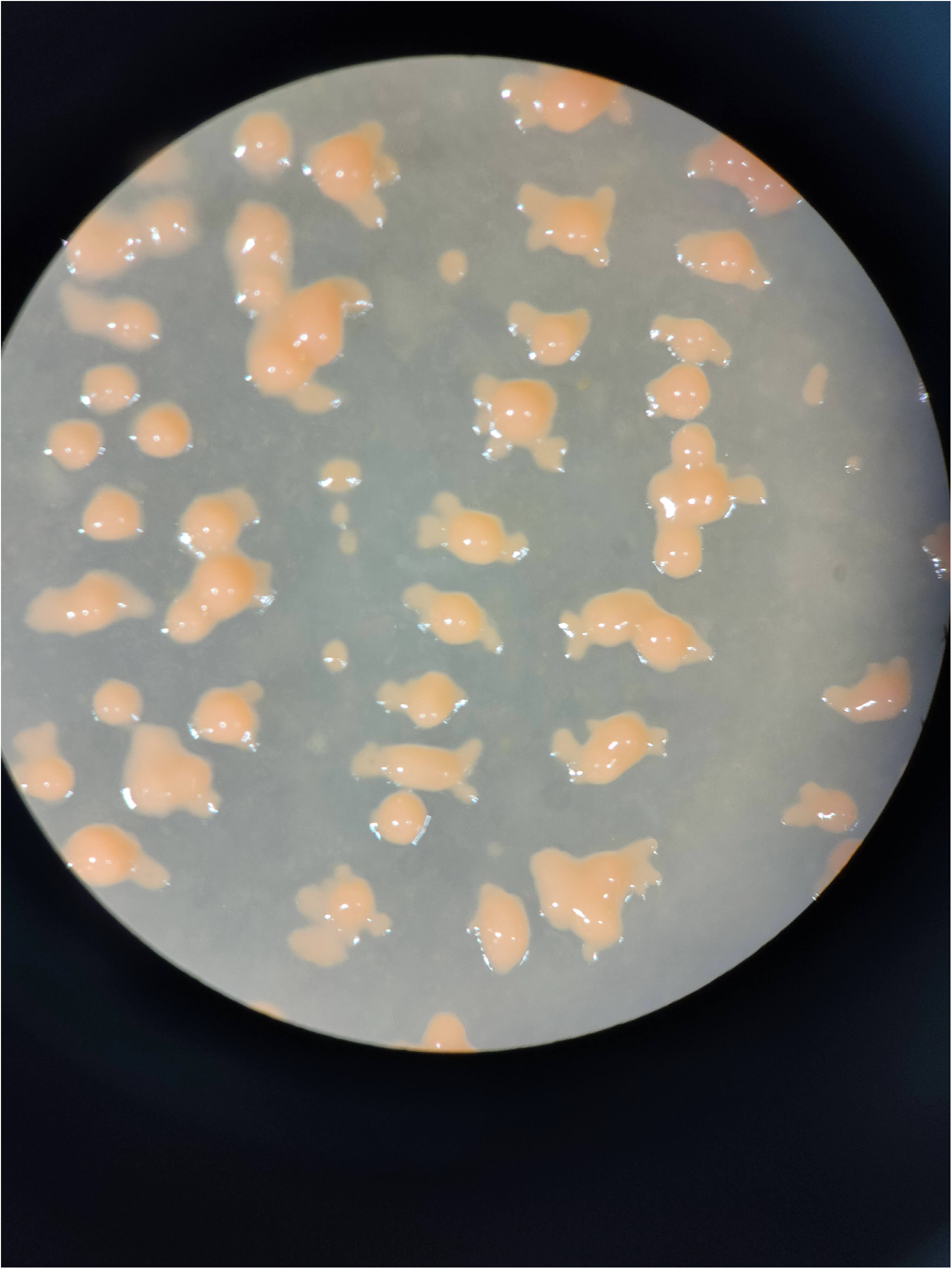

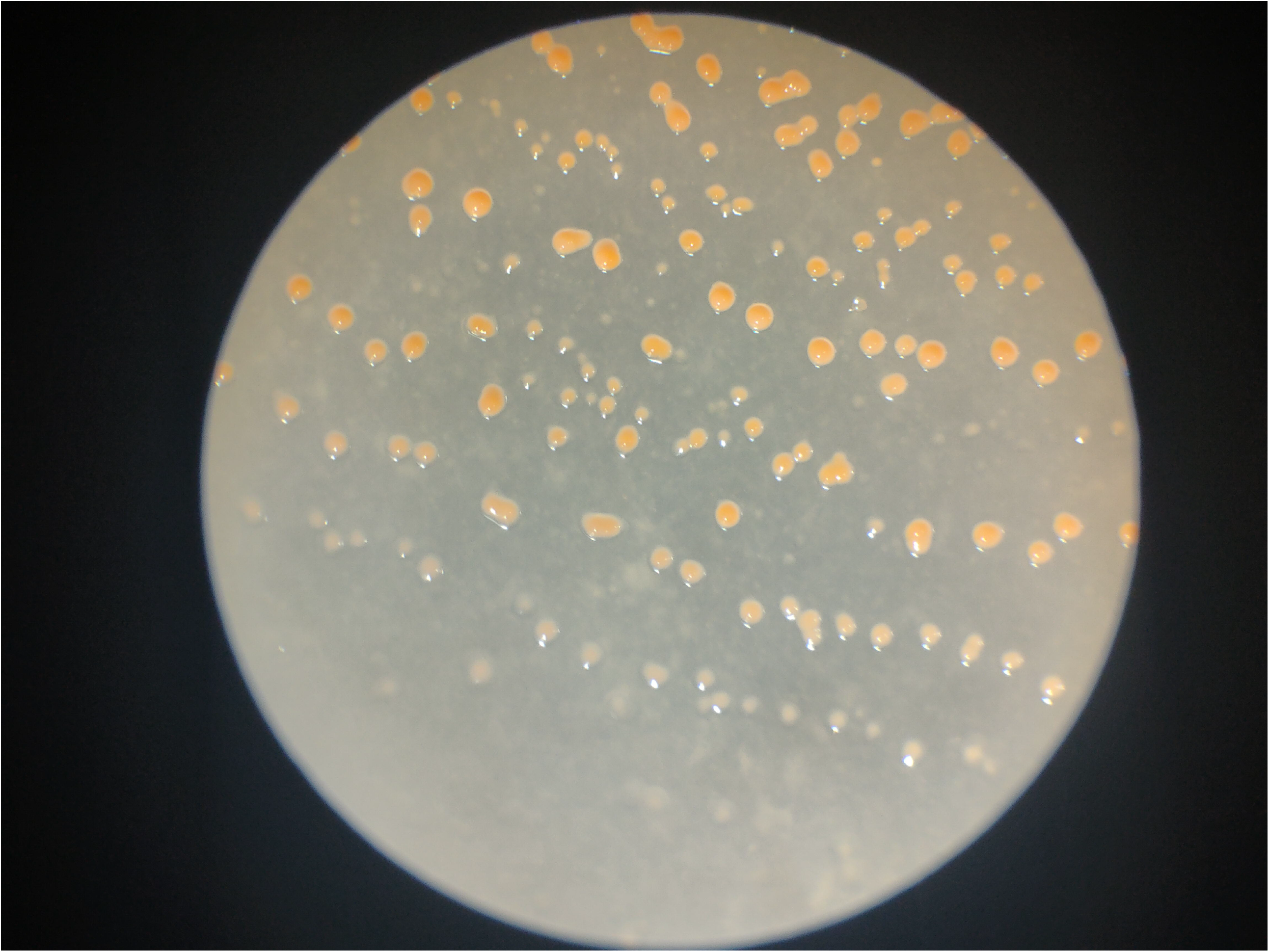

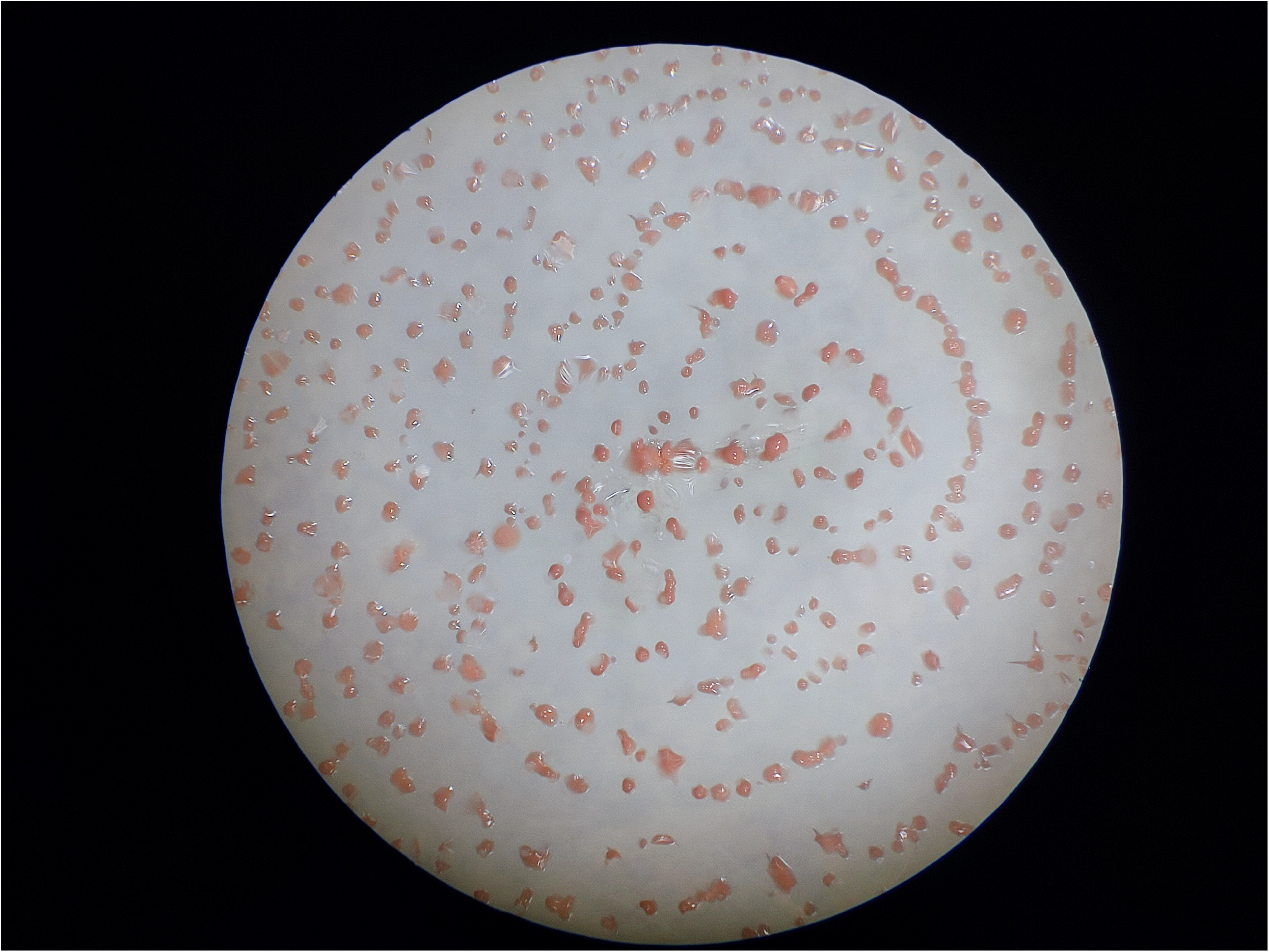

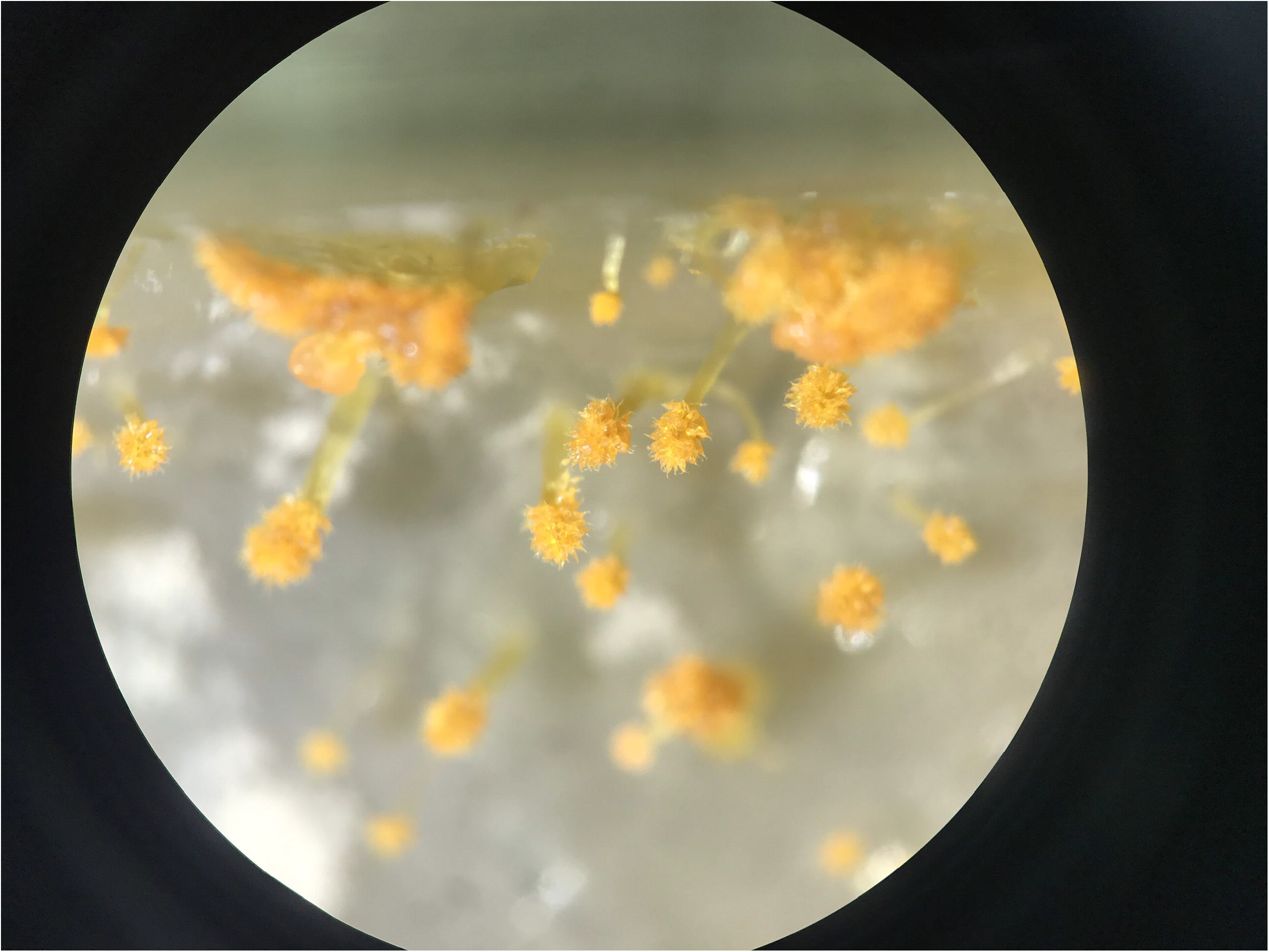

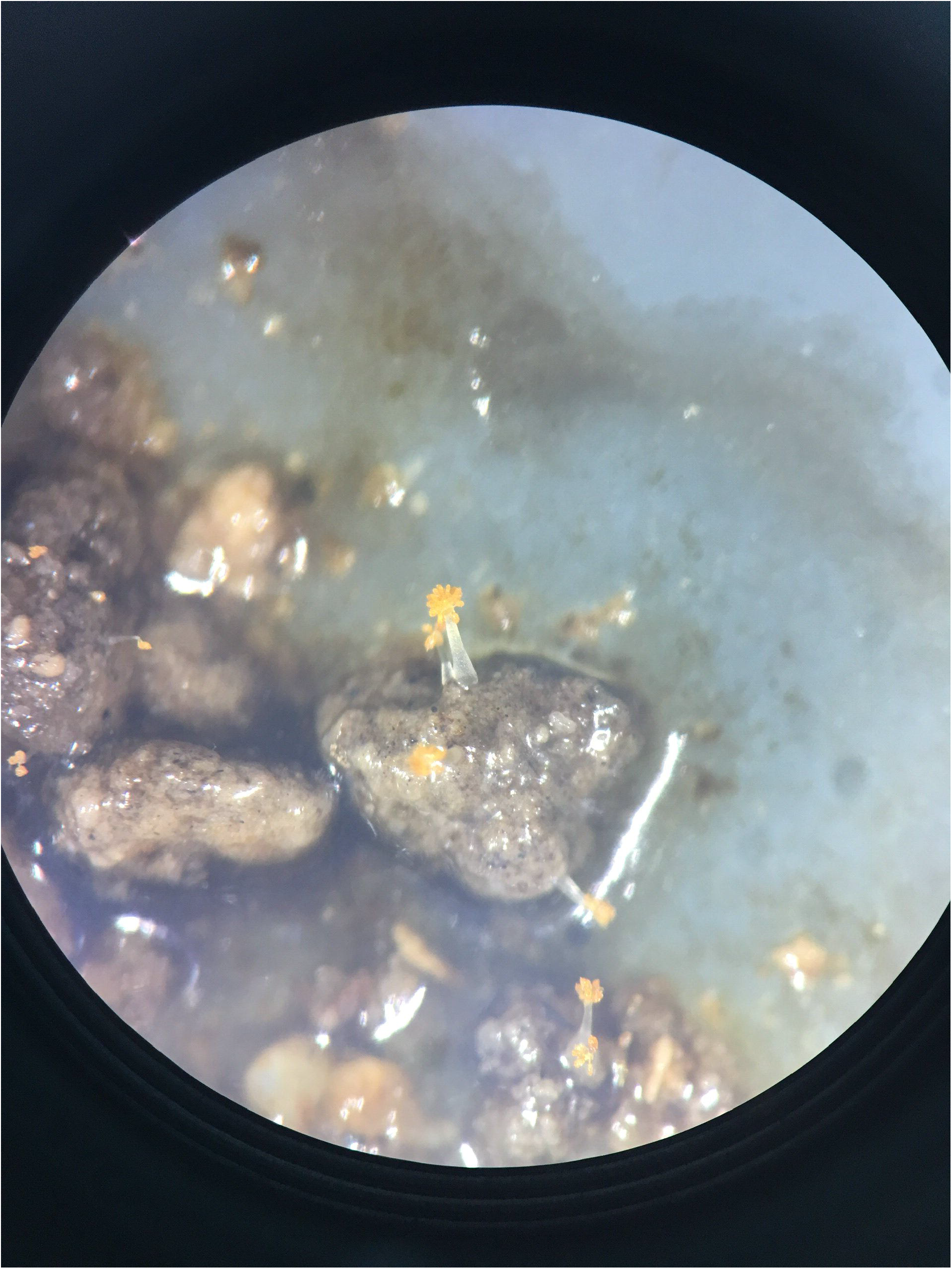

## Notes

### Competing Interest Statement

The authors have declared no competing interest.

