## Supplementary material for "MYXOBACTERIA FROM VIETNAM: ISOLATION, PHYLOGENETIC ANALYSIS AND BIOPROSPECTION": https://mail.google.com/mail/u/0/#inbox

Bảng 7.1. Kết quả thử nghiệm xác định $IC_{50}$ các chủng niêm khuẩn

| TT | Samples | DPPH assay | | ABTS assay | |
|---|---|---|---|---|---|
| | | $IC_{50}$ (µg/mL) | $IC_{50}$ ratio | $IC_{50}$ (µg/mL) | $IC_{50}$ ratio |
| | Trolox | | | 7.67 ± 0.82 | - |
| | Ascorbic acid | 10.90 ± 0.89 | - | | |
| 1 | AG11 | 116,27 ± 3,66 | 10,67 | 100,55 ± 1,13 | 13,11 |
| 2 | BD2 | 103,39 ± 2,90 | 9,49 | 128,33 ± 1,64 | 16,73 |
| 3 | BDi22 | 102.45 ± 2.75 | 9.40 | 65.25 ± 0.86 | 8.51 |
| 4 | BDi23 | 103,31 ± 2,70 | 9,49 | 135,05 ± 1,28 | 17,61 |
| 5 | BP161 | 172,41 ± 5,65 | 15,82 | 102,46 ± 1,24 | 13,36 |
| 6 | BP181 | 116,87 ± 3,73 | 10,73 | 117,05 ± 1,43 | 15,26 |
| 7 | BP182 | 90,39 ± 2,40 | 8,30 | 149,68 ± 2,27 | 19,51 |
| 8 | BP213 | 177,07 ± 5,16 | 16,25 | 128,53 ± 1,61 | 16,75 |
| 9 | BRVT1 | 123,63 ± 3,17 | 11,35 | 77,84 ± 0,92 | 10,15 |
| 10 | BT43 | 87.32 ± 2.35 | 8.01 | 85.02 ± 1.12 | 10.08 |
| 11 | BT92 | 96.35 ± 2.51 | 8.84 | 118.67 ± 1.67 | 15.47 |
| 12 | BT10 | 138,75 ± 3,70 | 12,73 | 102,74 ± 1,18 | 13,39 |
| 13 | CT21 | 52.34 ± 1.47 | 4.80 | 30.28 ± 0.74 | 3.95 |
| 14 | DN11 | 118,40 ± 3,59 | 10,87 | 119,28 ± 1,65 | 15,55 |
| 15 | DN21 | 130,63 ± 3,48 | 11,99 | 93,26 ± 1,12 | 12,16 |
| 16 | DN23 | 121,09 ± 2,98 | 11,11 | 92,74 ± 1,06 | 12,09 |
| 17 | GL41 | 57.24 ± 1.52 | 5.25 | 42.76 ± 0.50 | 5.57 |
| 18 | GL43 | 181,44 ± 5,34 | 16,65 | 120,29 ± 1,43 | 15,68 |
| 19 | HCM21 | 101,99 ± 2,61 | 9,36 | 87,50 ± 1,00 | 11,41 |
| 20 | HCM81 | 122.87 ± 3.35 | 11.28 | 67.14 ± 0.85 | 8.75 |
| 21 | HG223 | 121.30 ± 3.22 | 11.13 | 75.59 ± 0.94 | 9.85 |
| 22 | HG225 | 97,94 ± 2,57 | 8,99 | 119,04 ± 1,49 | 15,52 |
| 23 | LA41 | 149,99 ± 5,34 | 13,77 | 106,57 ± 1,23 | 13,89 |
| 24 | LA43 | 114,45 ± 3,08 | 10,50 | 97,18 ± 1,15 | 12,67 |
| 25 | LA44 | 81.38 ± 2.28 | 7.47 | 110.75 ± 1.35 | 14.44 |
| 26 | NB71 | 108,31 ± 2,87 | 9,94 | 130,10 ± 1,74 | 16,96 |
| 27 | NB8x1 | 169,96 ± 5,43 | 15,60 | 108,33 ± 1,33 | 14,12 |
| 28 | NB83 | 185,61 ± 6,05 | 1,04 | 91,37 ± 1,02 | 11,91 |
| 29 | PY1 | 140,27 ± 3,71 | 12,87 | 109,44 ± 1,24 | 14,27 |
| 30 | QT15 | 249,43 ± 6,17 | 22,89 | 188,81 ± 2,10 | 24,61 |
| 31 | QT72 | 167,33 ± 4,71 | 15,36 | 123,27 ± 1,50 | 16,07 |

| TT | Samples | DPPH assay | | ABTS assay | |
|---|---|---|---|---|---|
| | | $IC_{50}$ (µg/mL) | $IC_{50}$ ratio | $IC_{50}$ (µg/mL) | $IC_{50}$ ratio |
| 32 | ST11 | 182,90 ± 5,97 | 16,79 | 99,29 ± 1,13 | 12,94 |
| 33 | TG23 | 139,27 ± 4,66 | 12,78 | 93,77 ± 1,05 | 12,22 |
| 34 | TG131 | 40.28 ± 1.13 | 3.70 | 48.35 ± 0.58 | 6.30 |
| 35 | TH41 | 27.39 ± 1.74 | 2.51 | 116.70 ± 1.43 | 15.29 |
| 36 | TH48 | 144,13 ± 4,86 | 12,23 | 107,48 ± 1,27 | 14,01 |
| 37 | TH52 | 119,72 ± 2,96 | 10,99 | 87,04 ± 1,00 | 11,35 |
| 38 | TV11 | 164,24 ± 4,79 | 15,07 | 120,96 ± 1,45 | 15,77 |
| 39 | TV22 | 112,14 ± 2,88 | 10,29 | 133,00 ± 1,77 | 17,34 |
| 40 | TV51 | 116.13 ± 2.80 | 10.65 | 70.97 ± 0.86 | 9.25 |
| 41 | VL21 | 237,98 ± 5,69 | 21,84 | 197,36 ± 2,22 | 25,73 |
| 42 | VL25 | 136,18 ± 3,46 | 12,50 | 103,86 ± 1,18 | 13,54 |
| 43 | VL32 | 50.87 ± 1.33 | 4.67 | 57.22 ± 0.69 | 7.46 |

Bảng 7.2. Kết quả xác định MIC cao chiết các chủng niêm khuẩn

| Isolates | MR | MS | Sf | Ec | Pa | Ca | Rh | Mu | Pe | An |
|---|---|---|---|---|---|---|---|---|---|---|
| AG11 | 32 | 128 | - | - | - | - | - | 512 | 64 | 32 |
| BD2 | 4 | 4 | 32 | - | 512 | 8 | 4 | 8 | 2 | 1 |
| BDi22 | 128 | 512 | - | - | - | 16 | 32 | 64 | 1 | 1 |
| BDi23 | 32 | 8 | 128 | - | - | 512 | 256 | - | 512 | 64 |
| BP161 | 4 | 1 | 16 | - | 512 | 128 | 64 | 128 | - | - |
| BP181 | 1 | 1 | 4 | - | 512 | 64 | 128 | 256 | - | - |
| BP182 | 16 | 8 | 64 | - | 512 | 128 | 128 | 256 | - | - |
| BP213 | 32 | - | - | - | - | - | 512 | 512 | - | 512 |
| BRVT1 | - | - | - | - | - | - | 512 | - | 256 | 32 |
| BT43 | 16 | 8 | 128 | - | - | 128 | 256 | 512 | 16 | 4 |
| BT92 | 16 | 8 | 128 | - | 512 | 256 | - | 256 | 32 | 32 |
| BT10 | 64 | - | - | - | - | - | 256 | 256 | 256 | 512 |
| CT21 | 128 | 64 | 512 | - | - | 8 | 16 | - | 2 | 1 |
| DN11 | 32 | - | - | - | - | - | 512 | 512 | - | - |
| DN21 | 32 | 16 | 64 | - | 512 | 512 | 256 | 256 | 128 | 32 |
| DN23 | 512 | - | - | - | - | - | 512 | - | 512 | 1 |
| GL41 | 1 | 1 | 1 | 64 | 128 | 1 | 16 | 16 | 8 | 1 |
| GL43 | 32 | - | - | - | - | - | 256 | - | - | 512 |
| HCM21 | 32 | 128 | 512 | - | 512 | - | 128 | 256 | 256 | 64 |

| Isolates | MR | MS | Sf | Ec | Pa | Ca | Rh | Mu | Pe | An |
|---|---|---|---|---|---|---|---|---|---|---|
| HCM81 | 128 | - | - | - | 512 | 512 | 64 | 128 | - | - |
| HG223 | 64 | - | - | - | - | - | 256 | 512 | 512 | 512 |
| HG225 | 16 | 64 | 64 | - | 512 | 32 | 16 | 16 | 1 | 1 |
| LA41 | 64 | - | - | - | - | - | - | - | 512 | 512 |
| LA43 | 32 | 8 | 128 | - | 512 | 512 | 256 | 256 | 128 | 32 |
| LA44 | 64 | 16 | 256 | - | 512 | 512 | 256 | 256 | 1 | 32 |
| NB71 | 4 | 1 | 16 | - | 512 | 16 | 32 | 16 | 1 | 1 |
| NB8x1 | 64 | 16 | 256 | - | - | 512 | 64 | 64 | - | - |
| NB83 | 32 | - | - | - | - | - | - | - | 128 | 32 |
| PY1 | 64 | - | - | - | - | - | 256 | 256 | - | 512 |
| QT15 | 512 | 128 | 256 | - | - | 256 | - | - | - | - |
| QT72 | 32 | - | - | - | - | - | 128 | 256 | - | 512 |
| ST11 | 64 | - | - | - | - | - | 512 | - | 16 | 32 |
| TG23 | 64 | - | - | - | - | - | 512 | 256 | - | 512 |
| TG131 | 64 | - | - | - | - | - | 256 | 256 | 64 | 64 |
| TH41 | 128 | - | - | - | 512 | - | 256 | 256 | 256 | 32 |
| TH48 | 64 | 16 | 256 | - | - | 512 | 256 | 256 | 32 | 8 |
| TH52 | 128 | - | - | - | 512 | - | 256 | 256 | 512 | 256 |
| TV11 | 512 | 512 | - | - | - | - | 256 | 512 | - | 512 |
| TV22 | 64 | 128 | - | - | 512 | 512 | 128 | 256 | 32 | 32 |
| TV51 | 32 | - | - | - | - | 512 | 256 | - | 512 | 128 |
| VL21 | - | - | - | - | - | 512 | - | - | 128 | 128 |
| VL25 | - | - | - | - | - | 64 | 256 | 128 | 64 | 8 |
| VL32 | 64 | 256 | - | - | - | 512 | 128 | 64 | 1 | 4 |

Note: MR: MRSA, MS: MSSA, Sf: *S. faecalis*, Ec: *E. coli*, Pa: *P. aeruginosa*, Ca: *C. albicans*, Mu: Mucor sp., Rh: Rhizopus sp., Pe: Penicillium sp., and An: *A. niger*
